# SynGAP splice isoforms differentially regulate synaptic plasticity and dendritic development

**DOI:** 10.1101/2020.01.28.922013

**Authors:** Yoichi Araki, Ingie Hong, Timothy R. Gamache, Shaowen Ju, Leonardo Collado-Torres, Joo Heon Shin, Richard L. Huganir

## Abstract

SynGAP is a synaptic Ras GTPase-activating protein (GAP) with four C-terminal splice variants: α1, α2, β, and γ. Although recent studies have implicated *SYNGAP1* haploinsufficiency in ID/ASD pathogenesis, the degree to which each SynGAP isoform contributes to disease pathogenesis remains elusive. Here we demonstrate that individual SynGAP isoforms exhibit unique spatiotemporal expression and have distinct roles in neuronal and synaptic development. The SynGAP-α1 isoform, which undergoes robust liquid-liquid phase-separation with PSD-95 and is highly-enriched in synapses, is expressed late in development and disperses from synaptic spines in response to LTP-inducing synaptic activity to allow for AMPA receptor insertion and spine enlargement. In contrast, the SynGAP-β isoform, which undergoes less liquid-liquid phase-separation with PSD95 and is less synaptically targeted, is expressed early in development and promotes dendritic arborization. Interestingly, a SynGAP-α1 mutation that disrupts phase separation and synaptic targeting abolishes its function in plasticity and instead drives dendritic arbor development like the β isoform. These results demonstrate that distinct phase separation and synaptic targeting properties of SynGAP isoforms determine their function.

**Highlights:**

1. SynGAP-α1, α2, β, γ isoforms have distinct spatiotemporal expression and function in the brain.
2. SynGAP-α1 is required for plasticity, while β is required for dendritic development.
3. Liquid-liquid phase separation of SynGAP-α1 is required for its role in plasticity.
4. SynGAP isoforms may differentially contribute to SYNGAP1 related human NDDs.

## Introduction

SynGAP is a GTPase-activating protein (GAP) that is highly enriched in dendritic spines of excitatory neurons (Chen et al., 1998; Kim et al., 1998). SynGAP is a Ras- and Rap-GTPase activating protein that facilitates the hydrolysis of small G protein-bound GTP (active) to GDP (inactive), thus negatively regulating the activity of small G proteins (Carlisle et al., 2008; Chen et al., 1998; Pena et al., 2008; Rumbaugh et al., 2006). SynGAP is encoded by the *SYNGAP1* gene and is alternatively spliced to generate 4 distinct C-terminal isoforms: SynGAP-α1, SynGAP-α2, SynGAP-β, and SynGAP-γ (Li et al., 2001; McMahon et al., 2012). The C-terminal domain of SynGAP-α1 contains a class I PDZ ligand sequence (QTRV), which binds MAGUK family proteins such as PSD-95 (Chen et al., 1998; Kim et al., 1998; (Grant and O’Dell, 2001). Heterozygous deletion of *Syngap1* in rodents causes severe deficits in long-term potentiation (LTP) at synapses of hippocampal CA1 pyramidal neurons that are innervated by Schaffer collaterals (SC), as well as severe working memory deficits (Kim et al., 2003; Komiyama et al., 2002; Rumbaugh et al., 2006).

In humans, loss-of-function variants in *SYNGAP1* have been associated with Intellectual Disability (ID), epilepsy, Autism Spectrum Disorders (ASDs), and Neurodevelopmental Disability (NDD). While there are hundreds of genetic risk factors for these disorders, the significantly elevated frequency and 100% penetrance of loss-of-function variants in *SYNGAP1* as well as the range of brain disorders associated with *SYNGAP1* pathogenicity make it unique (Berryer et al., 2013; Carvill et al., 2013; Hamdan et al., 2011; Hamdan et al., 2009; Satterstrom et al., 2020).

Many loss-of-function variants of the *SYNGAP1* gene have been causally associated with ID, epilepsy, ASD, and other NDDs. In a UK study of 931 children with ID, *SYNGAP1* was the 4th most highly prevalent NDD-associated gene, and *SYNGAP1* variants accounted for ∼0.75 % of all NDD cases (UK-DDD-study, 2015). Patients with *SYNGAP1* haploinsufficiency have high rates of comorbid epilepsy, seizures, and acquired microcephaly (Berryer et al., 2013; Berryer et al., 2012; Carvill et al., 2013; Cook, 2011; Hamdan et al., 2011; Hamdan et al., 2009; Parker et al., 2015; Rauch et al., 2012; Tan et al., 2015; UK-DDD-study, 2015; Vissers et al., 2010; Vlaskamp et al., 2019; Writzl and Knegt, 2013). Mental Retardation, Autosomal Dominant 5 (MRD5) (OMIM #612621) is caused by mutations in *SYNGAP1.* MRD5 is characterized by moderate-to-severe intellectual disability with delayed psychomotor development apparent in the first years of life (Holder et al., 2019). Nearly all reported cases of *SYNGAP1*-related ID and ASD are *de novo* mutations within/near exons or splice sites of *SYNGAP1* (Vlaskamp et al., 2019).

Some key pathophysiological symptoms of ID and ASD observed in *SYNGAP1* patients have been recapitulated in constitutive *Syngap1* hetereozygous (*Syngap1*^+/-^) mice (Clement et al., 2012). *Syngap1* heterozygous mice exhibit learning deficits, hyperactivity and epileptic seizures (Clement et al., 2012; Guo et al., 2009). Additionally, several MRD5-associated *SYNGAP1* missense mutations also cause SynGAP protein instability (Berryer et al., 2013). These data strongly suggest that *SYNGAP1* haploinsufficiency is pathogenic in *SYNGAP1*-associated ID and ASD. Thus, several lines of evidence in mice and humans support that SynGAP is a critical regulator of synaptic plasticity, development and behavior.

We recently discovered that SynGAP-α1 is rapidly dispersed from dendritic spines during LTP, which allows for concomitant spine enlargement and accumulation of synaptic AMPARs (Araki et al., 2015). SynGAP-α1 dispersion from the dendritic spines releases the inhibition of synaptic RAS activity which is required for the expression of LTP (Harvey et al., 2008; Murakoshi and Yasuda, 2012; Walkup et al., 2016; Zhu et al., 2002). Additionally, SynGAP is the third mostly highly expressed protein in the postsynaptic density (PSD) and can undergo multivalent interactions with PSD-95 via liquid-liquid phase separation (LLPS), a process of forming highly concentrated condensates with liquid-like propensities, which may contribute to the formation of the PSD complex (Zeng et al., 2016). LLPS in cells is a phenomenon in which biochemical reactants are spatially clustered and concentrated in the absence of a surrounding membrane, allowing for organelle-like function without the physical and energetic barriers posed by lipid bilayers (Shin and Brangwynne, 2017). Although SynGAP is an ideal candidate to provide the structural basis of PSD (Zeng et al., 2018; Zeng et al., 2016), the phase separation of SynGAP was extensively characterized only with SynGAP-α1. The degree to which the other SynGAP isoforms undergo activity-dependent dispersion and LLPS, as well as the functional significance of these isoforms, remains largely unclear.

Although *SYNGAP1* haploinsufficiency likely affects the expression of all SynGAP isoforms, only the-α1 isoform has been rigorously characterized to date. Only a few functional studies of non-α1 SynGAP isoforms have been conducted to probe how these isoforms regulate synaptic physiology and disease pathogenesis (Li et al., 2001; McMahon et al., 2012). In these overexpression studies, the various SynGAP isoforms have been shown to have differing – and even opposing – effects on synaptic transmission (McMahon et al., 2012), however, as these were overexpression experiments, endogenous SynGAP was intact in this study complicating interpretation of these results. It is currently unknown whether *SYNGAP1*-associated ID/ASD pathology is associated with select deficits of specific SynGAP isoforms that may underlie unique features of NDD.

Here we report that SynGAP-α1 constitutes only 25-35% of total SynGAP protein in the brain, underscoring the importance of characterizing how the C-terminal SynGAP splice variants contribute to neuronal and synaptic development that are associated with the pathogenesis of *SYNGAP1* haploinsufficiency. In developing neurons, the various SynGAP isoforms have striking differences in neuroanatomical and subcellular expression. We report that SynGAP-β is expressed earliest in development, and functions to promote dendritic arbor development. In contrast, SynGAP-α1 reaches peak expression later in development and is a key regulator of synaptic plasticity mechanisms such as AMPAR insertion and dendritic spine enlargement. Our findings implicate unique roles for select SynGAP isoforms in mediating different facets of neuronal function. Furthermore, we identify isoform-specific differences in biochemical liquid-liquid phase-separation with PSD-95, and show how these differences inform the role each isoform plays in regulating synaptic plasticity or dendritic structure. These results suggest that individual SynGAP isoforms mediate distinct, specialized regulation of neuronal and synaptic development and have implications for potential therapeutic treatments for *SynGAP1* related disorders.

## Results

### SynGAP isoforms have distinct and overlapping expression profiles during brain development

*SYNGAP1* is alternatively spliced at several sites to include exons 18, 19, or 20 to generate four unique C-terminal isoforms: SynGAP-α1, SynGAP-α2, SynGAP-β, and SynGAP-γ (**Fig. 1*A, B***). SynGAP-α1 and SynGAP-α2 isoforms skip exon 19 and are produced by selective splicing of exon 20, whereby SynGAP-α1 contains a PDZ ligand (-QTRV) and SynGAP-α2 lacks this domain. The SynGAP-β isoform includes a frameshifting extension of exon 18 leading to early termination, which generates a SynGAP protein with a partially truncated coiled-coil domain. The SynGAP-γ isoform includes exon 19, which contains a short coding sequence followed by a STOP codon (-LLIR*).

**Figure 1.**
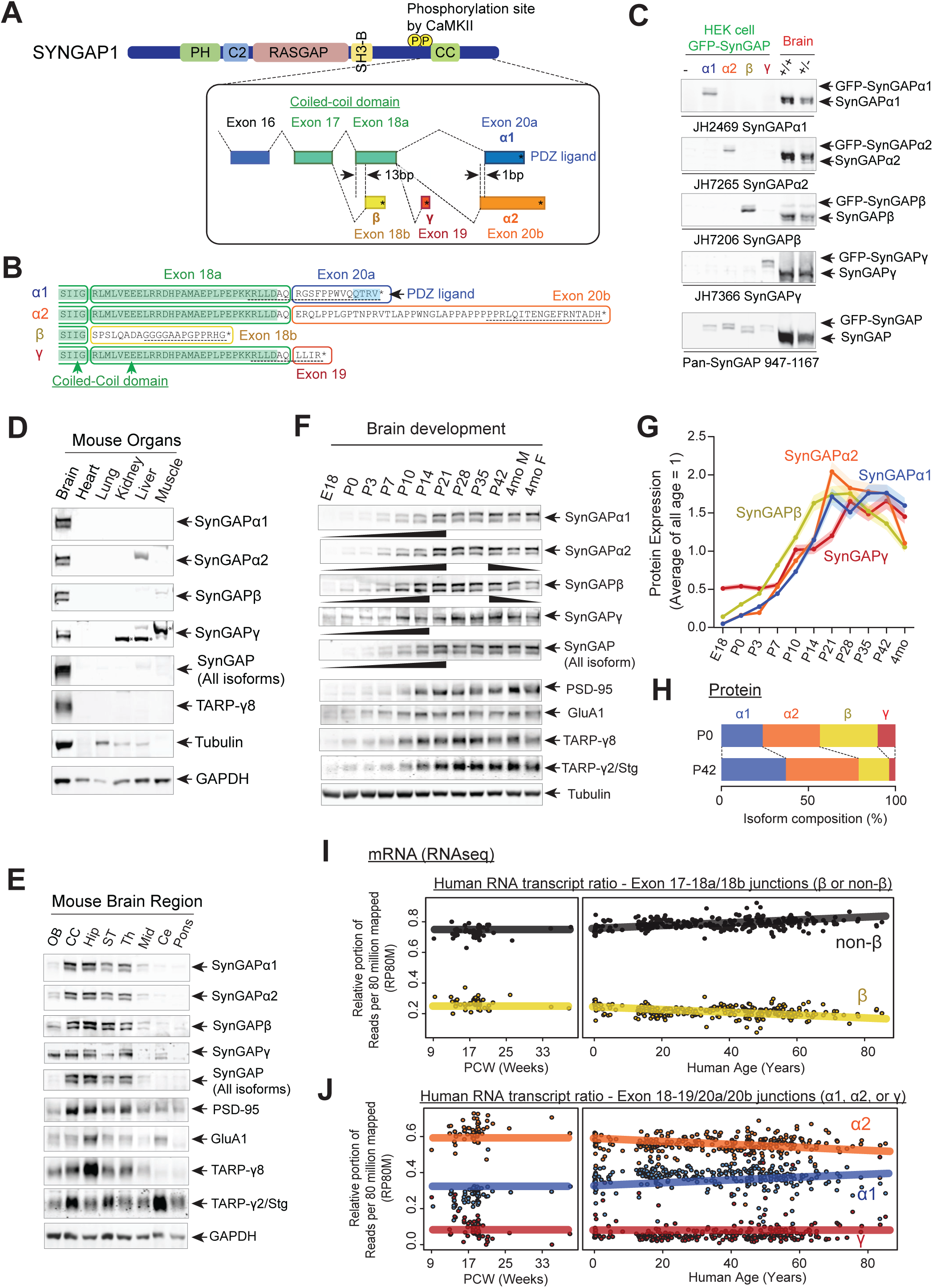
SynGAP isoforms are differentially expressed during brain development. (A) Schematic of *SYNGAP1* splicing at the C-terminus. *SYNGAP1* is alternatively spliced within exons 18-20 to generate four unique C-terminal isoforms designated as α1, α2, β, and γ. (B) C-terminal amino-acid sequences of SynGAP isoforms encoding select protein domains. Coil-Coil domain (yellow) and PDZ ligand-binding domain (blue). Targeted epitopes of isoform-specific SynGAP antibodies (JH2469, JH7265, JH7206, and JH7366) are indicated. (C) Specificity of SynGAP isoform-specific antibodies. Immunoblots of SynGAP isoform expression in lysates prepared from HEK 293T cells expressing individual GFP-tagged SynGAP isoforms and lysates prepared from brain tissue obtained from *WT* and *Syngap1* +/- mice were shown. Quantification of relative SynGAP isoform levels with respect to total SynGAP expression measured from immunoblot were shown in Supplementary Fig. 1A. Two-way ANOVA followed by Tukey’s post hoc test (Isoform F(4,30) = 1.900; p = 0.13, Genotype F(1,30) = 451.2; p < 0.001, Interaction F(4,30)=1.900; p = 0.13, n = 4 each condition) was performed. Error bar indicates ± SEM. (D) Endogenous expression and distribution of SynGAP isoforms in various organs. Immunoblots of qualitative distribution of SynGAP isoforms in lysates prepared from various organ tissues of WT mice were shown. Asterisks indicate non-specific bands that are also detected in tissue from knockout mice. Two-way ANOVA followed by Tukey’s post hoc test (Tissue F(5,144) = 1433; p < 0.0001, Isoform F(7,144) = 229.3; p < 0.0001, Interaction F(35,144) = 25.45; p<0.0001, n = 4 each condition) was performed. Heat map of immunoblots was displayed in Supplementary Fig. 1B. The amount of protein in the brain is standardized as 1.0. (E) Western blot of endogenous levels of individual SynGAP isoforms and other synaptic proteins in lysates prepared from several brain regions obtained from *WT* and *Syngap1* +/- mice. (OB: Olfactory bulb, CC: Cerebral cortex, Hip: Hippocampus, ST: Striatum, Th: Thalamus, Mid: Midbrain, Ce: Cerebellum). Two-way ANOVA followed by Tukey’s post hoc test (Brain regions F(7, 264) = 1048; p < 0.0001, Molecules F(10,264) = 8.0 × 10 ^-12^; p > 0.9999, Interaction F(70.264) = 59.06; p<0.0001, n = 4 each condition) was performed. Graph showing the mean values of each signals was displayed in Supplementary Fig. 1C. (F-H) Developmental expression profiles of individual SynGAP isoforms and related synaptic proteins. (F) Immunoblots of SynGAP isoform expression measured in forebrain tissue lysates prepared from *WT* and *Syngap1* +/- mice at different developmental ages. (G) Quantification of immunoblots representing relative enrichments along developmental stage. The mean values of each signals were plotted in the graph. (H) Quantification of absolute SynGAP isoform abundance at P0 and P42 from Fig. 1C and 1G. Error bars indicate ± SEM. Two-way ANOVA followed by Tukey’s post hoc test (Developmental stage F(10,330) = 397.4; p<0.0001, Molecule F(9,330) = 2.116; p = 0.027, Interaction F(90,330) = 26.18; p < 0.0001, n = 4 each condition) was performed. (I) mRNA expression of the β and non-β SYNGAP1 isoforms across age in human dorsolateral prefrontal cortex. The relative portion of RNAseq reads spanning the exon 17-18 junction supporting either isoform was plotted against human age (post-conception weeks and years) with a linear regression. (J) mRNA expression of the α1, α2, and γ SYNGAP1 isoforms across age. The relative portion of RNAseq reads spanning the exon 18-19 junction (γ) or 18-20 (α1, α2) junction supporting each isoform was plotted against human age. Reads per 80 million mapped (RP80M) of RNAseq data were shown in Supplementary Fig. 1E, F.

To characterize each SynGAP isoform, we raised antibodies using SynGAP C-terminal peptides as antigens (**Fig. 1*B***, black underlines). Antibody specificity was validated in transfected HEK cells (**Fig. 1*C*** Left 4 lanes, and quantification in **Supplementary Fig. 1*A***), as well as in brain lysate from WT and *Syngap1* heterozygote (*Syngap1 +/-*) mice, in which immunoblotting demonstrates an expected ∼50% reduction of expression of all SynGAP isoforms (**Fig. 1*C*** Right 2 lanes, quantification in **Supplementary Fig. 1*A***). All four SynGAP isoforms are enriched in brain tissue (* asterisks: non-specific band) with other brain-specific proteins, such as Stargazin and TARP-γ8 (**Fig. 1*D***, and quantification in **Supplementary Fig. 1*B***). To determine the expression profile of SynGAP isoforms, we isolated 8 brain regions from adult (P42) mice. All four SynGAP isoforms are enriched in forebrain regions such as the cerebral cortex and hippocampus in comparison to hindbrain structures such as the pons (**Fig. 1*E***, quantification in **Supplementary Fig. 1*C***). However, there are several isoform-specific differences in regional expression. For example, SynGAP-β and SynGAP-γ are also expressed in the olfactory bulb, and SynGAP-γ is expressed at a significant level in the cerebellum. *SYNGAP1* mutations have been linked to NDDs such as ID and ASD, which suggests an important role for *SYNGAP1* in normal brain development. Thus, we sought to investigate the expression of the SynGAP isoforms throughout development in brain tissue from mice at several developmental stages spanning late embryogenesis to adulthood (**Fig. 1*F-H***, complete set of quantification in **Supplementary Fig. 1*D***). SynGAP-β is well expressed early in development (E18-P14) whereas SynGAP-α2 is generally the most abundant isoform and reaches maximal abundance at P21-P35. SynGAP-α1 expression increases later in development. (**Fig. 1*F-H***). Expression of other synaptic proteins (GluA1, PSD-95, and TARPs) reached maximum between P21 and P42, which is similar to the timeframe for maximal expression of SynGAP-α1 and SynGAP-α2.

In order to more rigorously quantify the expression levels of SynGAP isoforms over development, using standardized detection ratios of each isoform to total SynGAP based on Figure 1C, we calculated the relative abundance (% total SynGAP) of each isoform at P0 and P42 (**Fig. 1*F-H***). SynGAP-β is relatively highly expressed at P0 (34.6 ± 0.6 %) and decreases to 15.7 ± 0.8 % at P42. SynGAP-α2 increases slower than the β isoform prenatally but is also expressed well at P0 (31.9 ± 0.4 %). The α2 isoform is the most abundant isoform at the P42 (44.9 ± 1.5 %), which is consistent with a previous finding that SynGAP-α2 is the dominant isoform at the mRNA level as well (Yokoi et al., 2017). SynGAP-α1 exhibits relatively low expression levels at P0 (24.3 ± 0.3 %) and then accumulates throughout development, eventually becoming the second most highly-expressed isoform when measured at P42 (35.0 ± 0.9 %) next to the α2 isoform. SynGAP-γ is expressed at low levels throughout development (9.1± 0.5 % at P0, and 4.3 ± 0.3 % at P42) (**Fig. 1*H***).

To extend our protein-level observations and investigate the correspondence to human SYNGAP1, we analyzed previously published human brain RNAseq data (n = 338) (Jaffe et al., 2018) (Fig. 1*I,J*, Reads per 80 million mapped (RP80M) in **Supplementary Fig. 1*E, F***). Consistent with our biochemical estimates, splice junctions that lead to the β isoform comprised ∼22% of all reads spanning the exon 17-18 junction and decreased slightly across development (**Fig. 1*G***). At the α1/ α2/γ junction, which is relevant in the non-β transcripts (the remaining 78%), we observe that junction reads corresponding to the α2 isoform are the most abundant across all ages (∼56% of reads spanning exon 18 to 19/20 junctions), and the α1 isoform follows at ∼35%, increasing slowly throughout development, while the γ isoform junction reads are rare (∼9% of non-β) (**Fig. 1*J***). This correspondence with protein data shows that the SynGAP isoform abundance levels are tightly controlled on the level of splicing and the ratios across development are conserved in mice and humans. Therefore, we next tested whether these isoforms have unique neuronal functions and play distinct roles in SYNGAP1-related pathogenesis.

### Unique LLPS propensities of SynGAP isoforms correlate with their localization to the cytoplasm or the post-synaptic density

To better understand isoform-specific function in neurons, we first investigated differences in liquid-liquid phase separation (LLPS), a mechanism for the effective subcellular organization of cellular proteins. We previously discovered that SynGAP-α1 undergoes LLPS with PSD-95 at physiological concentrations *in vitro*, resulting in the concentration of SynGAP into dense condensates that are reminiscent of the postsynaptic density (Zeng et al., 2016). To investigate the biochemical and phase separation propensities of other SynGAP isoforms, we first performed a sedimentation assay in HEK cells transfected with constructs encoding tagged full-length PSD-95 and SynGAP (**Fig. 2*A***). Here, centrifugation of the sample resulted in two fractions: the phase-separated, insoluble-protein-containing pellet fraction (termed: **[P])** and the soluble supernatant fraction (termed: **[S])**. The ratio of each protein in the condensed phase fraction (**[P]** /(**[S]** + **[P]**), termed “Phase-separation index”) was calculated to indicate the propensity of the protein to undergo LLPS. Both myc-PSD-95 and GFP-SynGAP WT remain mostly in the soluble fraction when expressed alone in HEK cells (23.1 ± 4.2 % of PSD-95 in the condensed phase fraction, 38.2 ± 0.5 % of SynGAP-α1 in the condensed phase fraction when expressed alone, Supplementary Fig 2A). Co-expression of both myc-PSD-95 and GFP-SynGAP-α1 WT causes a dramatic increase in their concentration in the phase-separated [P] fraction (80.3 ± 2.2 % of PSD-95 and 74.7 ± 3.3 % of SynGAP-α1 in the condensed phase fraction when co-expressed, *** p < 0.001 compared to expressed alone). We previously generated a mutant form of SynGAP-α1 that contains two point mutations in SynGAP-α1 – L1202D and K1252D (GFP-SynGAP-α1 LDKD) (Zeng et al., 2016) – which prevents SynGAP trimerization and phase-separation with PSD-95. Consistent with the hypothesis that the coil-coiled domain of SynGAP-α1 is critical for LLPS with PSD-95 co-sedimentation of SynGAP-α1 LDKD with PSD-95 was significantly decreased in the [P] fraction when compared to that of GFP-SynGAP-α1 WT with PSD-95 (44.0 ± 5.0 % of PSD-95 and 27.3 ± 4.6 % of SynGAP-α1 LDKD at condensed phase fraction when SynGAP-α1 LDKD and PSD-95 was co-expressed, *** p < 0.001 compared to SynGAP-α1 WT and PSD-95 was co-expressed, **Supplementary Fig 2*A***). We also determined the PDZ ligand (QTRV) of SynGAP-α1 is critical for efficient phase separations in this assay (**Supplementary Fig. 2*B***, Phase separation index: 80.0 ± 3.8 % of PSD-95 when co-expressed with SynGAP-α1, but 49.7 ± 2.7 % of PSD-95 when co-expressed with SynGAP-α1 ΔQTRV, ** p< 0.01). These data are consistent with the results of *in vitro,* cell-free sedimentation assay experiments reported previously (Zeng et al., 2016).

**Figure 2.**
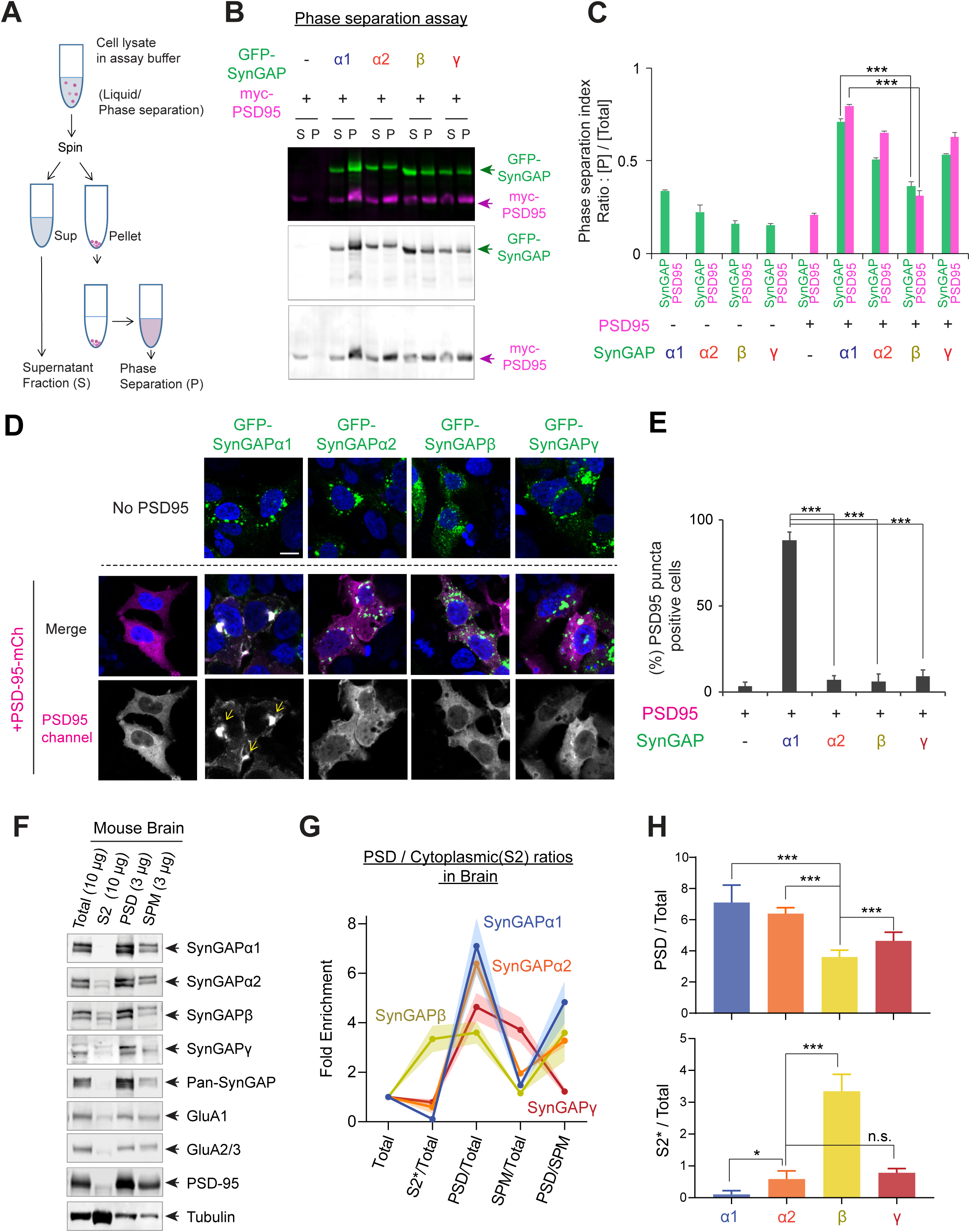
Phase separation and subcellular localization of various SynGAP isoforms. (A) Schematic diagram of LLPS sedimentation assay. HEK 293T cell lysates were centrifuged and fractionated into insoluble pellet (P) and soluble supernatant (S) fractions. (B) Representative western blot probing levels of individual SynGAP isoforms in phase-separated supernatant and pellet lysate fractions obtained from HEK cells expressing myc-PSD95 and individual GFP-tagged SynGAP isoforms. (C) Quantification of pellet fraction ratios obtained from averaged western blots as the representative example shown in (D). Error bars indicate ± SEM. Two-way ANOVA followed by Tukey’s post hoc test (Transfections F(8,54) = 812,2; p < 0.0001, Molecules F (1,54) = 50.88; p < 0.0001, Interaction F(8,54) = 101.5; p < 0.0001, n = 4, *** p<0.001, ** p<0.01, * < 0.05) was performed. (D, E) Representative confocal images of living HEK cells expressing myc-PSD95 alone or myc-PSD95 and individual SynGAP isoforms. Scale Bar, 10 μm (D). (E) Quantification of the averaged percentage of PSD95-positive puncta identified in images of living HEK cells as shown in (D). Error bars indicate ± SEM. One-way ANOVA ANOVA followed by Tukey’s post hoc test (Transfections F (4, 15) = 96.77; p < 0.0001, n = 4 independent coverslip, *** p<0.001, ** p<0.01, * < 0.05) was performed. (F-H) Immunoblot probing endogenous levels of individual SynGAP isoforms and other synaptic proteins in forebrain tissue lysates obtained from adult mice subjected to postsynaptic density fractionation. (G, H) Averaged enrichment of SynGAP isoforms in subcellular fractions in comparison to their levels within the total homogenate fraction, S2 fractions, and PSD fractions. Error bars indicate ± SEM. Two-way ANOVA followed by Tukey’s post hoc test (Fractions F(4,145) = 274.6; p < 0.0001, Molecules F (7,145) = 77.13; p < 0.0001, Interaction F(28,145) = 206.7; p < 0.0001, n = 4-7 independent samples for each molecules, *** p<0.001, ** p<0.01, * < 0.05) was performed.

We next examined the propensity of each SynGAP isoform to undergo LLPS with PSD-95 (**Fig. 2*B, C***, and **Supplementary Fig 2*C***). Expressed separately, all individual isoforms were preferentially found in the soluble fraction. Co-expression of individual SynGAP isoforms with PSD-95 dramatically increased the presence of both GFP-SynGAP-α1 and myc-PSD-95 in the phase-separated fraction (71.0 ± 1.4 % of SynGAP-α1 in condensed phase fraction when co-expressed with PSD-95). Similarly, GFP-SynGAP-α2 and GFP-SynGAP-γ also exhibited an increased localization in pellet fraction, albeit to a lesser extent than that of GFP-SynGAP-α1 (50.8 ± 0.9 % of SynGAP-α2 and 53.3 ± 0.5 % of SynGAP-γ in condensed phase fraction when co-expressed with PSD-95, **Fig. 2*B, C***, and **Supplementary Fig 2*D***) These isoforms harbor a complete coiled-coil domain but lack the PDZ ligand. In contrast, GFP-SynGAP-β and myc-PSD-95 did not efficiently co-sediment (36.2 ± 2.5 % of SynGAP-β in condensed phase fraction when co-expressed with PSD-95. *** p < 0.001, compared to SynGAP-α1 coexpressed with PSD-95). SynGAP-β lacks the PDZ ligand and contains only a partial coiled-coil domain. These results highlight the necessity and additive functions of the coiled-coil domain and PDZ ligand in mediating SynGAP isoform interactions and binding to PSD-95.

We next used confocal microscopy to assess SynGAP isoform-dependent biomolecular condensate formation in living HEK cells (**Supplementary Fig 2*D***). We previously reported that GFP-SynGAP-α1 and RFP-PSD-95 undergo LLPS in living cells and form liquid-like cytoplasmic droplets when expressed in living cells (Zeng et al., 2016). When expressed alone in HEK 293T cells PSD95-mCherry (PSD95-mCh) exhibited relatively diffuse cytoplasmic expression (4.8 ± 1.0 % of PSD-95 puncta positive cells). In contrast, co-expression of PSD95-mCh and GFP-SynGAP-α1 WT led to a dramatic increase in distinct cytoplasmic puncta (> 1 μm diameter) (85.5 ± 3.3 % of PSD-95 puncta positive cells when co-expressed, *** p < 0.001 compared to PSD-95 alone). However, GFP-SynGAP-α1 LDKD did not induce puncta formation when co-expressed with PSD-95-mCh (22.3 ± 5.3% of PSD-95 puncta positive cells, *** p < 0.001 compared to SynGAP-α1 WT and PSD-95 was co-expressed) (**Supplementary Fig. 2*D***). We next determined the percentage of cytoplasmic puncta-positive cells following co-expression of PSD-95-mCh along with each SynGAP isoform. SynGAP-α1 expression robustly induced distinct puncta with PSD-95 (88.2 ± 4.9 % of PSD-95 puncta positive cells when SynGAP-α1 and PSD-95 were coexpressed) that were absent with expression of all other SynGAP isoforms (7.3 ± 2.3 %, 6.0 ± 4.3%, 9.0 ± 3.9 % when SynGAP-α2, β,γ and PSD-95 were co-expressed respectively, *** p < 0.001, compared to SynGAP-α1 and PSD-95 co-expression) (**Fig. 2*D, E***). The failure of non-SynGAP-α1 isoforms to induce the formation of measurable cytoplasmic puncta suggests that a complete coiled-coil domain and PDZ-ligand are necessary for the LLPS of SynGAP in this assay. These results suggest that SynGAP isoforms have unique LLPS properties that are determined by their C-terminal sequences.

Finally, we examined biochemical distributions of SynGAP isoforms in the mouse brain (**Fig. 2*F-H***, complete set of quantification in **Supplementary Fig. 2*E***). Mouse brains were excised and fractionated into Total (Total homogenate), S2 (13,800 x g Supernatant), SPM (Synaptosomal plasma membrane), and PSD (Post-synaptic density). Almost SynGAP isoforms were highly enriched in PSD fractions (α1 7.1 ± 0.5 fold enrichment, α2 6.3 ± 0.2 fold enrichment, β 3.6 ± 0.2 fold enrichment, γ 4.6 ± 0.3 fold enrichment). However, the SynGAP-β isoform was significantly less enriched in PSD (*** p < 0.001, PSD enrichment of SynGAP-β compared to α1 and α2). Additionally, SynGAP-β was significantly more highly expressed in the S2 (cytosolic) fraction compared to other isoforms. In contrast, the expression of the α1 isoform was very low in this fraction ([S2]*10/[Total] ratio; β 0.33 ± 0.02, *** p < 0.001, compared to other isoforms α1 0.011 ± 0.005, α2 0.059 ± 0.011, γ 0.079 ± 0.006). These results indicate that phase separation characteristics of SynGAP isoforms *in vitro* reflects the *in vivo* biochemical localization of SynGAP isoforms.

### SynGAP isoforms differentially regulate GTPase activity to Ras, Rap1, and Rac1

Because the SynGAP isoforms are differentially expressed throughout the brain and possess distinct abilities to associate with synaptic scaffolds, we investigated if differences also exist among SynGAP isoforms in their ability to activate RAS family GTPases. To test this possibility, we assayed levels of GTP-bound GTPases such as Ras, Rap1, and Rac1 in HEK cells expressing several small G-proteins in the presence of individual SynGAP isoforms. Our data demonstrate that specific SynGAP isoforms differentially activate GTP hydrolysis bound to Ras, Rap1, and Rac1 (**Fig. 3*A-C***). SynGAP-β possesses the highest GAP activity among all isoforms (50.6 ± 3.7 % decrease in Ras-GTP, 53.3 ± 6.7 % decrease in Rap1-GTP, 39.2 ± 2.7 % decrease in Rac1-GTP) (**Fig. 3*D, E***). SynGAP-α1 preferentially activate GTP-hydrolysis of Ras (28.2 ± 2.8 % decrease in Ras-GTP), compared to Rap1 (14.3 ± 4.3 % decrease in Rap1-GTP, * p < 0.05 compared to Ras) (**Fig. 3*D, E***). SynGAP-α2 has a similar trend to decrease Ras-GTP over Rap1-GTP (37.0 ± 3.5 % decrease in Ras-GTP, compared to 24.8 ± 1.0 % decreases in Rap1-GTP, p = 0.06). Conversely, SynGAP-β highly activates GTP hydrolysis of Rap1 with similar levels with Ras (53.2 ± 6.8 % decreases in Rap1-GTP, compared to 50.6 ± 3.8 % decreases in Ras-GTP) (**Fig. 3*D,E***). Since small G proteins were not phase-separated, the most soluble SynGAP-β isoform may have the highest exposure to cytosolic small G-proteins, and thus, have the highest GAP activities. It is also possible that different C-terminal structures enhance or decrease the accessibility of various small G proteins to the GAP domain and thus differentially regulate their GAP activity.

**Figure 3.**
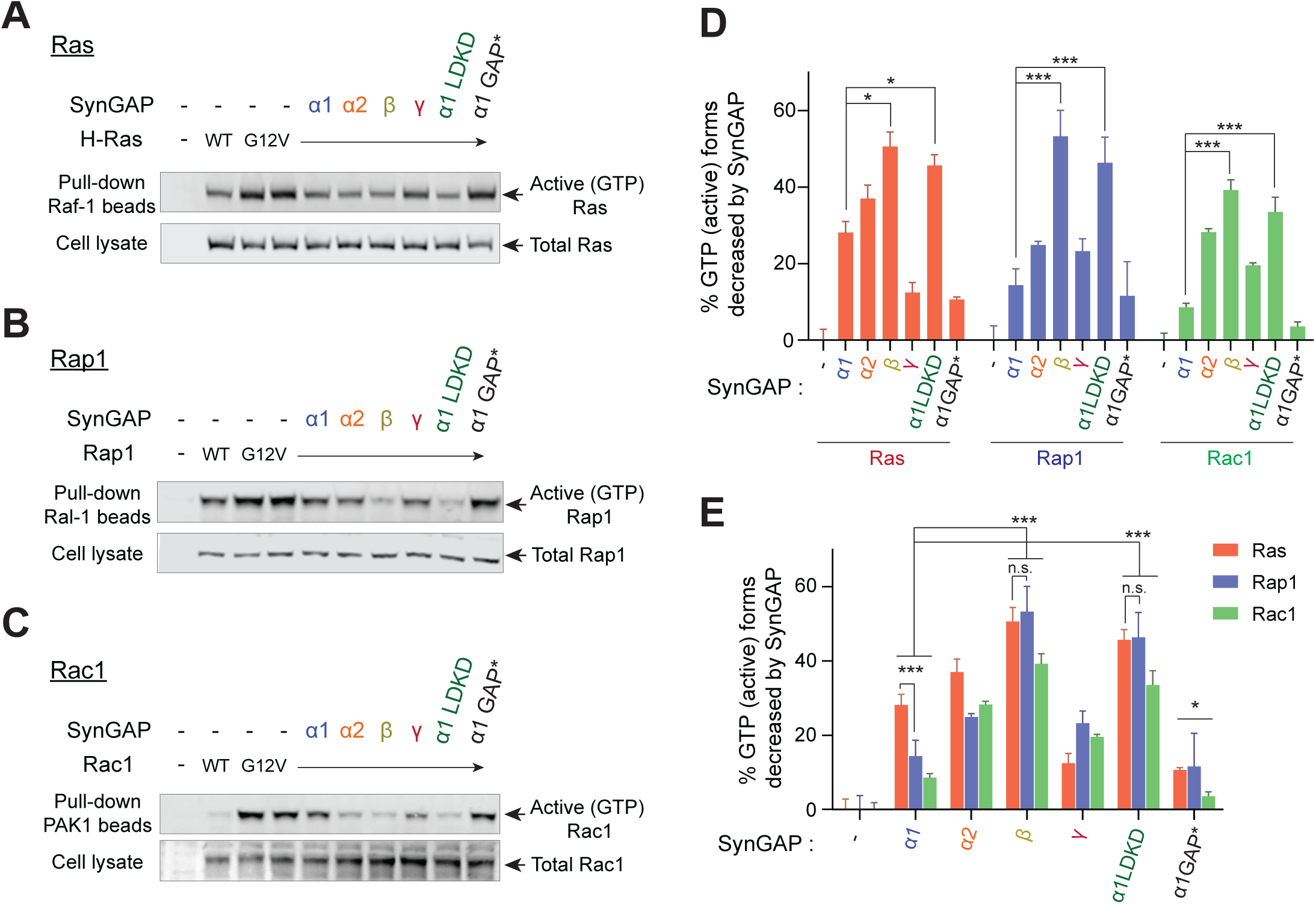
SynGAP isoforms differentially regulate the activity of small G proteins. (A) Representative immunoblot detecting levels of active GTP-bound Ras following co-immunoprecipitation of active Ras by pulldown of Raf1 in response to expression of individual SynGAP isoforms in HEK cell lysates. (B) Representative immunoblot detecting levels of active GTP-bound Rap1 following co-immunoprecipitation of active Rap1 by pulldown of Ral1 in response to expression of individual SynGAP isoforms in HEK cell lysates. (C) Representative immunoblot detecting levels of active GTP-bound Rac1 following co-immunoprecipitation of active Rac1 by pulldown of PAK1 in response to expression of individual SynGAP isoforms in HEK cell lysates. (D, E) Quantification of averaged percent reduction of active GTP-bound forms of Ras, Rap1, and Rac1 normalized to total (active + inactive) levels in response to expression of individual SynGAP isoforms expressed in HEK cell lysates. Error bars indicate ± SEM. Two-way ANOVA followed by Tukey’s post hoc test (SynGAP isoforms F(6,105) = 62.76; p < 0.0001, small G proteins F (2,105) = 7.414; p = 0.0010, Interaction F(12,105) = 2.207; p = 0.016, n = 6, *** p<0.001, ** p<0.01, * < 0.05) was performed.

### Differential dispersion dynamics of SynGAP isoforms during LTP

Previously, we have shown that SynGAP-α1 undergoes rapid NMDAR-CaMKII-dependent dispersion from the synapse, which is required for AMPAR insertion and spine enlargement during LTP (Araki et al., 2015). In order to investigate the dispersion dynamics of the other SynGAP isoforms during LTP, we employed a knockdown-replacement strategy in cultured hippocampal neurons, whereby endogenous SynGAP expression was depleted via shRNA-mediated knockdown and individual GFP-tagged, shRNA-resistant SynGAP isoforms were transfected (**Fig. 4**). We knocked down 77.3 % ± 0.1 % of endogenous SynGAP by shRNA and replaced with similar amount (−90% of endogenous proteins) by shRNA resistant SynGAP isoform construct (**Supplementary Fig. 3**). Cultured neurons were subjected to a chemical LTP (chemLTP) treatment during live confocal imaging, and the amount of synaptically localized GFP-tagged SynGAP as well as dendritic spine size were measured before and after LTP (**Fig. 4*A, B***). In this chemLTP stimulation, the magnesium in the media was withdrawn in conjunction with glycine perfusion. With spontaneous glutamate release from axonal terminals, glycine strongly and specifically stimulates synaptic NMDA receptors (Liao et al., 2001; Lu et al., 2001). SynGAP-α1 exhibited high synaptic localization prior to LTP induction and then underwent rapid dispersion following LTP (3.5 ± 1.3 fold synaptic spine enrichment of SynGAP-α1 before chemLTP, 1.7 ± 0.3 fold synaptic spine enrichment after chemLTP, *** p < 0.001) (**Fig. 4*A, B***). GFP-SynGAP-α2 was also dispersed albeit to a lesser extent than GFP-SynGAP-α1 (2.8 ± 0.6 fold synaptic spine enrichment of SynGAP-α2 before chemLTP, 1.8 ± 0.4 fold synaptic spine enrichment after chemLTP, * p < 0.05). In contrast, GFP-SynGAP-β is less enriched at synapses and fails to disperse upon chemLTP stimulation (1.9 ± 0.09 fold synaptic spine enrichment of SynGAP-α2 before chemLTP, 1.4 ± 0.11 fold synaptic spine enrichment after chemLTP, not significant p > 0.05). (**Fig. 4*A, B***). Taken together, these data indicate distinct dynamics for individual SynGAP isoforms such that SynGAP-α1 was most robustly dispersed during LTP and the β isoform was not.

**Figure 4.**
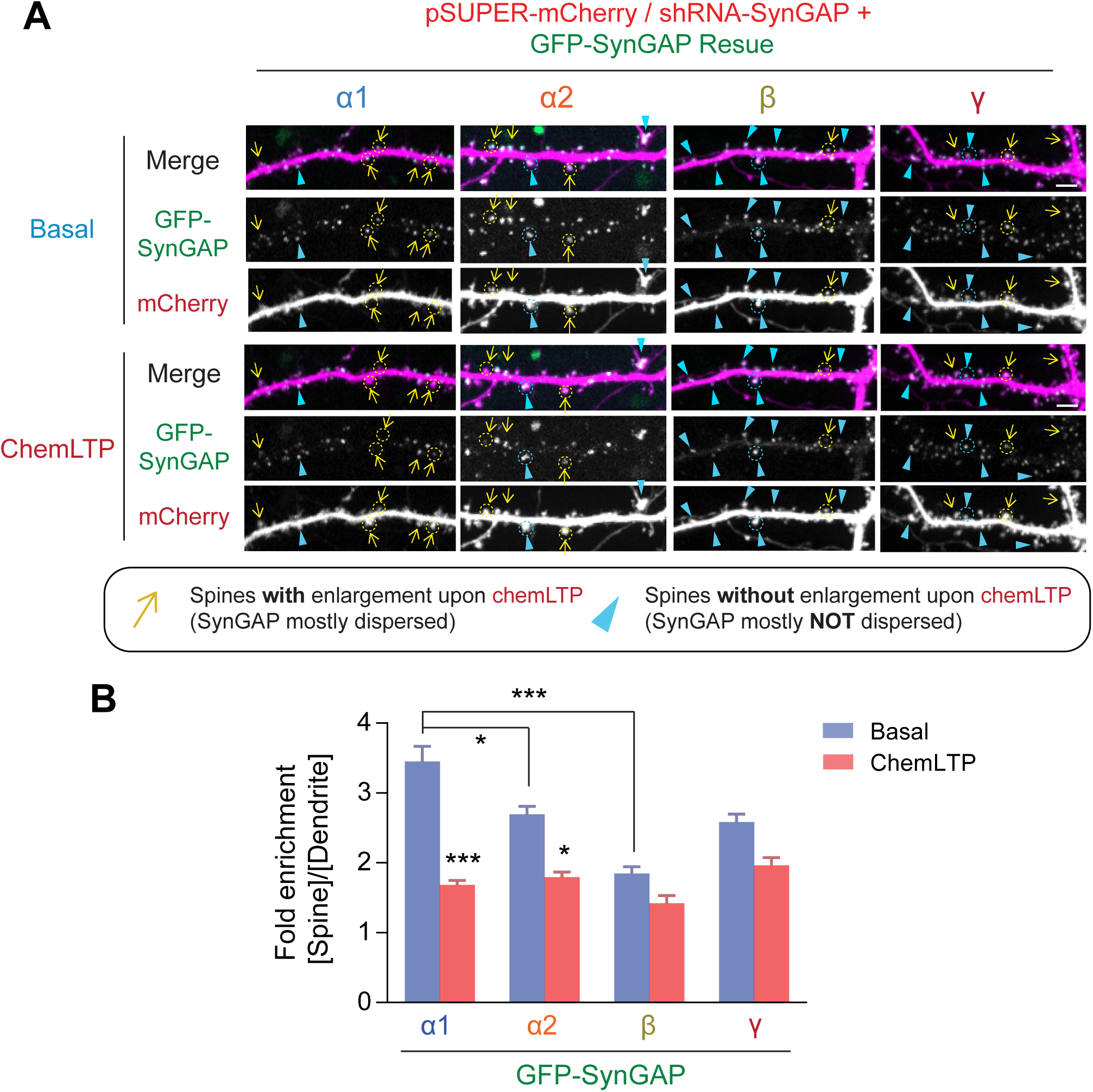
Dispersion dynamics of various SynGAP isoforms from synaptic spines during LTP. (A) Live confocal images of hippocampal neurons expressing individual GFP-tagged SynGAP isoforms and mCherry during basal conditions and chemLTP conditions. Yellow arrows indicate dendritic spines that enlarged following chemical LTP treatment. Blue arrowheads mark dendritic spines that did not enlarge after chemical LTP treatment. Scale Bar, 5 μm. (B) Quantification of averaged relative change in the GFP-SynGAP isoform at synaptic spines following chemLTP treatment. Synaptic localization of SynGAP isoforms was determined by calculating the ratio of GFP intensity within dendritic spine heads and dividing by GFP intensity localized to the dendritic shaft at the base of the dendritic spine. Error bars indicate ± SEM. Two-way ANOVA followed by Tukey’s post hoc test (SynGAP isoforms F(3,296) = 21.43; p < 0.0001, chemLTP F (1,296) = 119.9; p < 0.0001, Interaction F(3,296) = 6.607; p < 0.0001, n = 37 - 39 spines from 4 independent experiments, *** p<0.001, ** p<0.01, * < 0.05) was performed. Error bar indicates ± SEM.

### Synaptic AMPAR insertion and spine enlargement during LTP are differentially regulated by SynGAP isoforms

We previously demonstrated that SynGAP-α1 undergoes rapid NMDAR- and CaMKII-dependent dispersion from the synapse, and this dispersion is required for synaptic AMPAR insertion and spine enlargement that occur during LTP (Araki et al., 2015). So far, we have determined that various SynGAP isoforms have distinct phase separation propensities, have differential GAP activity, synaptic dendritic localization, and dispersion kinetics during LTP. Thus, we hypothesize these SynGAP isoforms function differentially during LTP. To test this hypothesis, we replaced endogenous SynGAP with shRNA-resistant Azurite-tagged SynGAP isoforms (Araki et al., 2015; Zeng et al., 2016). Transfected neurons also expressed the pH-sensitive Super ecliptic pHluorin (SEP)-tagged-GluA1 (SEP-GluA1) and mCherry to monitor surface AMPAR expression and dendritic spine size in response to chemLTP treatment (**Fig. 5*A-E***) (Lin et al., 2009). Under control conditions, significant increases in synaptic membrane-localized AMPARs and dendritic spine size were observed following LTP stimulation (2.5 ± 1.2 fold synaptic enrichment of AMPA receptor in synaptic spines [*** p < 0.001] and 2.7 ± 1.4 fold synaptic spine size [*** p < 0.001] after chemLTP compared to basal condition) (**Fig. 5*B*** and **Fig. 5*C-E***). Dendritic spine enlargement and synaptic AMPAR accumulation at synapses were occluded when endogenous SynGAP expression was depleted via shRNA-mediated knockdown; this is due to elevated Ras activity, spine enlargement and synaptic AMPAR accumulation in unstimulated baseline conditions (Araki et al., 2015) (2.0 ± 1.2 fold enrichment of AMPA receptor in basal state to 2.3 ± 0.9 fold enrichment after chemLTP [Not significant, p > 0.05] / 2.1 ± 1.6 fold synaptic spine size in basal state to 2.5 ± 1.6 fold spine size after chemLTP [Not significant, p > 0.05] in *SYNGAP1*-shRNA) (**Fig. 5*A1*** and **Fig. 5*C-E***). Molecular replacement with SynGAP-α1 restored baseline SEP-GluA1 and mCherry intensities to levels comparable to those measured in baseline control conditions and rescued LTP-dependent enhancement of dendritic spine volume and surface AMPAR content (1.1 ± 0.7 fold enrichment of AMPA receptor in basal state to 2.3 ± 0.6 fold enrichment after chemLTP [*** p < 0.001] / 1.4 ± 0.8 fold synaptic spine size in basal state become 2.4 ± 0.7 fold spine size after chemLTP [*** p < 0.001] in *SYNGAP1*-shRNA + SynGAP-α1 expression) (**Fig. 5*A2*** and **Fig. 5*C-E***). SynGAP-α2 underwent modest dispersion following stimulation and rescued basal spine enlargement and AMPAR insertion after chemLTP to a much lesser extent compared to SynGAP-α1 (1.7 ± 1.3 fold enrichment of AMPA receptor in basal state to 2.3 ± 1.2 fold enrichment after chemLTP [* p < 0.05] / 1.8 ± 0.8 fold synaptic spine size in basal state become 2.5 ± 0.4 fold spine size after chemLTP [Not significant p = 0.06] in *SYNGAP1*-shRNA + SynGAP-α2 expression) (**Fig. 5*A3*** and **Fig. 5*C-E***). Replacement with SynGAP-β totally failed to rescue basal spine enlargement and AMPAR insertion following chemLTP treatment (2.0 ± 0.8 fold enrichment of AMPA receptor in basal state to 2.5 ± 0.8 fold enrichment after chemLTP [Not significant, p > 0.05] / 2.4 ± 1.1 fold synaptic spine size in basal state to 2.8 ± 1.8 fold spine size after chemLTP [Not significant, p > 0.05] in *SYNGAP1*-shRNA + SynGAP-β expression) (**Fig. 5*A4*** and **Fig. 5*C-E***). We previously found that the phase separation mutant of SynGAP-α1 (LDKD) only partially rescued the LTP and significantly lowered the LTP threshold (Zeng et al., 2016). These results suggest that both the coiled-coil domain and PDZ ligand are required for LTP rescue in SynGAP KD neurons, and only SynGAP-α1 harbors the necessary and sufficient domains for efficient LTP expression. Our data suggest a specialized role for SynGAP-α1 in regulating LTP.

**Figure 5.**
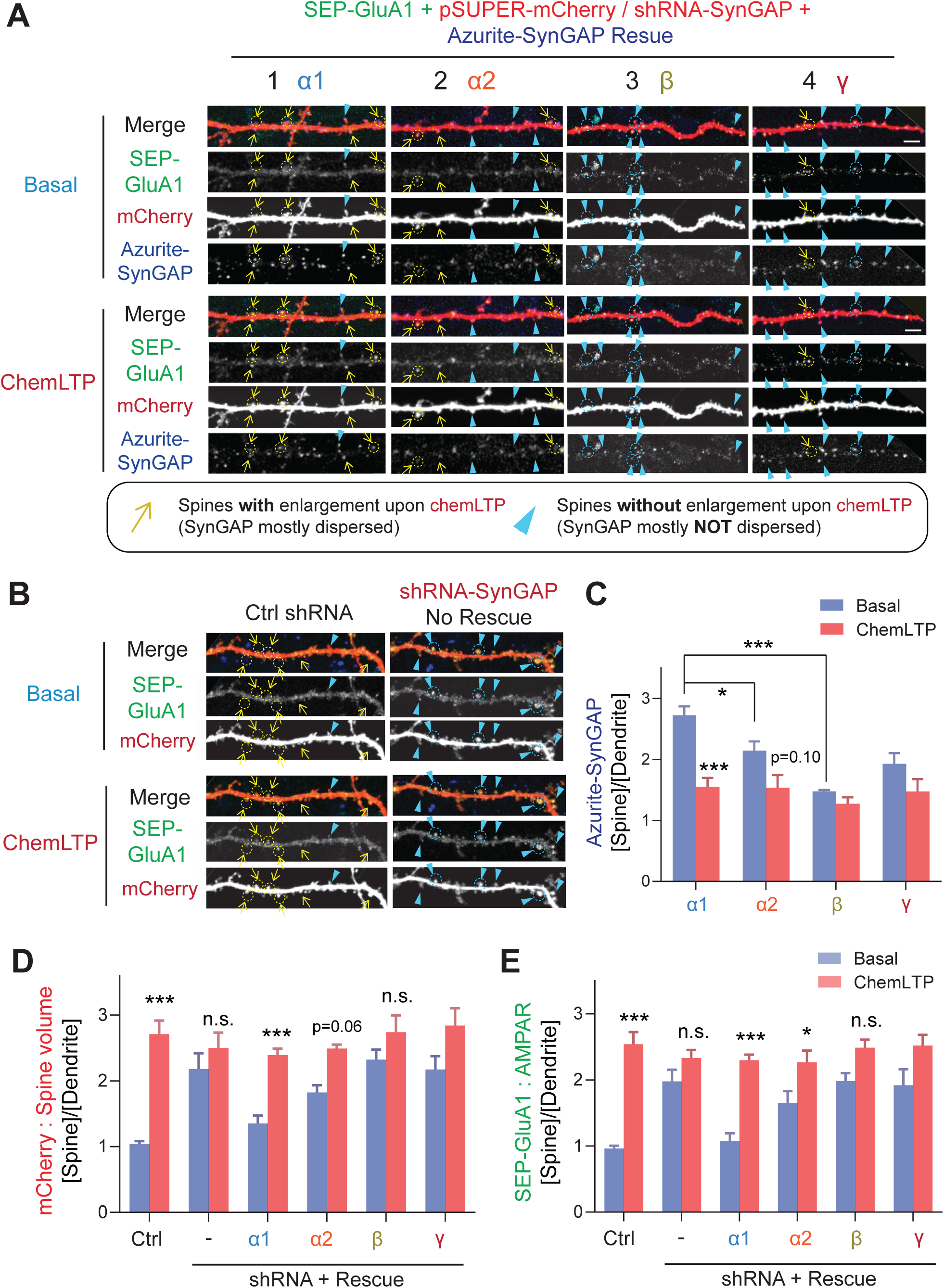
SynGAP-α1 rescues AMPA receptor trafficking and structural plasticity deficits in SynGAP-depleted hippocampal neurons. (A, B) Representative live confocal images of cultured hippocampal neurons expressing SEP-GluA1, mCherry, and individual Azurite-tagged SynGAP isoforms in basal and chemLTP conditions. Endogenous SynGAP was knocked-down and replaced with individual Azurite-tagged SynGAP isoforms. Yellow arrows indicate dendritic spines that exhibited LTP-induced enlargement. Scale Bar, 5 μm. (C-E) Quantification of averaged SEP-GluA1, mCherry, Azurite-SynGAP intensity in dendritic spines on hippocampal neurons expressing individual SynGAP isoforms before and after chemLTP treatment. Error bars indicate ± SEM. Two-way ANOVA followed by Tukey’s post hoc test (ChemLTP F(1,242) = 501.1 [GluA1], 426.4 [mCherry], 219.4 [SynGAP], p < 0.001; Genotype F(5,242) = 30.68 [GluA1], 35.71 [mCherry], 553.7 [SynGAP], P < 0.001; Interaction (5,240) = 15.02 [GluA1], 18.57 [mCherry], 553.7 [SynGAP], p < 0.001; n = 47-49 spines from 4 independent coverslips each condition, *** p<0.001, ** p<0.01, * < 0.05) was performed. Error bar indicates ± SEM.

### Dendritic arbor development is regulated by select SynGAP isoforms

Since our data reveal only modest roles for non-α1 isoforms in regulating synaptic plasticity in contrast to the differences in G-protein activity-regulation, we decided to investigate whether SynGAP isoforms are involved in other aspects of neuronal function. A previous report showed that *Syngap1* +/- mice have a significantly altered dendritic arborization pattern (Aceti et al., 2015). Thus, we assessed the effects of specific SynGAP isoforms in regulating dendritic development. In this experiment, the effects of *SYNGAP1* knockdown on dendritic branching was assessed by comparing control and *SYNGAP1* shRNA-expressing hippocampal neurons at DIV 8 (**Fig. 6*A***). Obvious basal (< 10-50 μm in length) dendrites (Control: 3.4 ± 0.5 intersections at 10 μm) and a branched primary apical (>100-150 μm in length) dendrites emanate from the somas of control neurons (Control: 4.2 ± 0.3 intersections at 150 μm). Sholl analysis revealed that *SYNGAP1* knockdown aberrantly enhances the number of neurite extensions proximal to neuronal cell bodies (8.3 ± 0.8 interactions at 10 μm, *** p < 0.001 compared to Control 3.4 ± 0.5 intersections at 10 μm) (**Fig. 6*B, F***). In contrast, *SYNGAP1* knockdown significantly decreased distal branches (*SYNGAP1*-shRNA : 0.3 ± 0.2 intersections at 150 μm, *** p < 0.001 compared to Control : 4.2 ± 0.3 intersections at 150 μm) (**Fig. 6*B, F***). The aberrantly elevated outgrowth of neurites proximal to neuron somas that is associated with *SYNGAP1* knockdown was rescued by expressing all SynGAP isoforms (α1 rescue : 4.0 ± 0.5 intersections at 10 μm, α2 rescue : 3.0 ± 0.3 intersections at 10 μm, β rescue : 3.7 ± 0.5 intersections at 10 μm, γ rescue 4.3 ± 0.2 intersections at 10 μm, *** p < 0.001 compared to *SYNGAP1*-shRNA : 8.3 ± 0.8 interactions at 10 μm) (**Fig. 6*C, G***). Interestingly, only SynGAP-β effectively rescued the distal dendritic complexity deficits (150 μm) by restoring the formation of primary and secondary apical dendrites (β rescue : 3.8 ± 0.6 intersections at 150 μm *** p < 0.001 compared to *SYNGAP1*-shRNA 0.3 ± 0.2 intersections at 150 μm). All other α1, α2, and γ isoforms failed to rescue distal branching deficits (α1 rescue : 1.7 ± 0.4 intersections at 150 μm, α2 rescue : 1.3 ± 0.2 intersections at 150 μm, γ rescue : 1.0 ± 0.3 intersections at 150 μm, not significant compared to *SYNGAP1*-shRNA : 0.3 ± 0.2 interactions at 150 μm) (**Fig. 6*C, G***). Interestingly, expression of SynGAP-α1 LDKD rescued the primary dendrite phenotype, similar to the effect of expression of SynGAP-β (α1 LDKD rescue : 3.5 ± 0.4 intersections at 150 μm *** p < 0.001, β rescue : 3.8 ± 0.6 intersections at 150 μm ***p < 0.001, compared to *SYNGAP1*-shRNA : 0.3 ± 0.2 intersections at 150 μm) (**Fig. 6*C, D, H***). This result suggests that disruption of SynGAP-α1 LLPS results in more β like behavior, rescuing distal dendritic arbor deficits. Finally, we found that a GAP mutant of SynGAP (Araki et al., 2015) does not rescue the dendritic arbor deficits (α1 GAP* rescue : 7.7 ± 1.2 intersections at 10 μm, not significant compared to *SYNGAP1*-shRNA : 8.3 ± 0.8 interactions at 10 μm), indicating that GAP activity is required for the dendritic phenotype rescue described here (**Fig. 6*E, F***).

**Figure 6.**
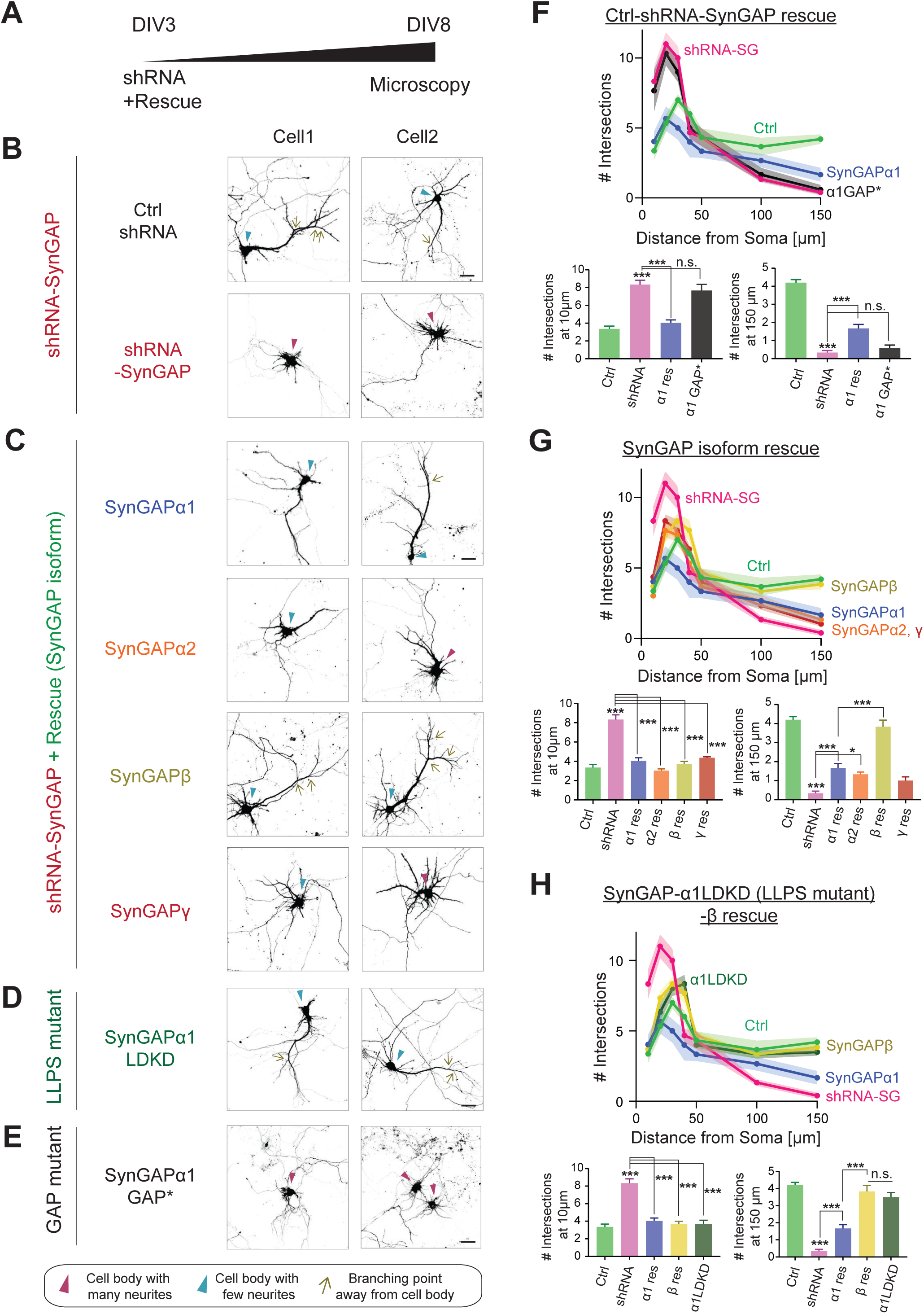
SynGAP-β rescues aberrant dendritic arbor development in SynGAP-depleted neurons. (A) Schematic of experimental timeline for assessing the effects of individual SynGAP isoform expression on dendritic development. Neuronal morphology was evaluated by observing co-transfected DsRed at DIV 8. (B-E) Representative images of dendritic arbors of young cultured hippocampal neurons expressing DsRed upon SynGAP knockdown (B) and upon *SYNGAP1* knockdown plus expressing individual SynGAP isoforms (C), SynGAPα1 LLPS mutant (D), or SynGAPα1 GAP mutant (E). Scale Bar, 20 μm. (F-H) Sholl analysis of dendritic branches presented as the mean number of intersections plotted as a function of distance from the center of the cell body (center = 0). Error bars indicate ± SEM. Two-way ANOVA followed by Tukey’s post hoc test (Distance F(6,952) = 288.6, p < 0.001; Genotype F(7,952) = 21.96, P < 0.001; Interaction (42,952) = 14.83, p < 0.001, n = 18) was performed. Error bar indicates ± SEM.

## Discussion

Here, we have characterized the developmental expression and subcellular localization, biochemical properties, and functional roles of each C-terminal SynGAP splice variant. Our data suggest novel roles for SynGAP isoforms in regulating dendrite and synapse development in neurons. SynGAP-β is expressed at higher portions early in brain development, and is gradually replaced in the mature brain by increasing expression of SynGAP-α1 and SynGAP-α2. Although SynGAP-β appears dispensable for synaptic plasticity function, it exhibits the strongest GAP activity of all the C-terminal isoforms, preferentially targets Rap1, and facilitates dendritic arbor development. SynGAP-β not only lacks a PDZ ligand, but also lacks a full coiled-coil domain, likely leading to the dramatically reduced LLPS propensity and increased cytoplasmic localization. SynGAP-α1, however, contains a complete coiled-coil domain and PDZ ligand, leading to the strongest LLPS and robust concentration at the PSD and is critical for LTP expression. This dense packing in the PSD in turn may be important to allow the dynamic dispersion of SynGAP during LTP (**Fig. 7*A***), which we have shown earlier to be required for both spine growth and AMPAR trafficking (Araki et al., 2015). The LLPS mutant of SynGAP-α1 failed to rescue synaptic plasticity but instead rescued dendritic arbor development like SynGAP-β, suggesting that LLPS propensity determines the roles of SynGAP in the regulation of neuronal maturation and/or synaptic plasticity.

**Figure 7.**
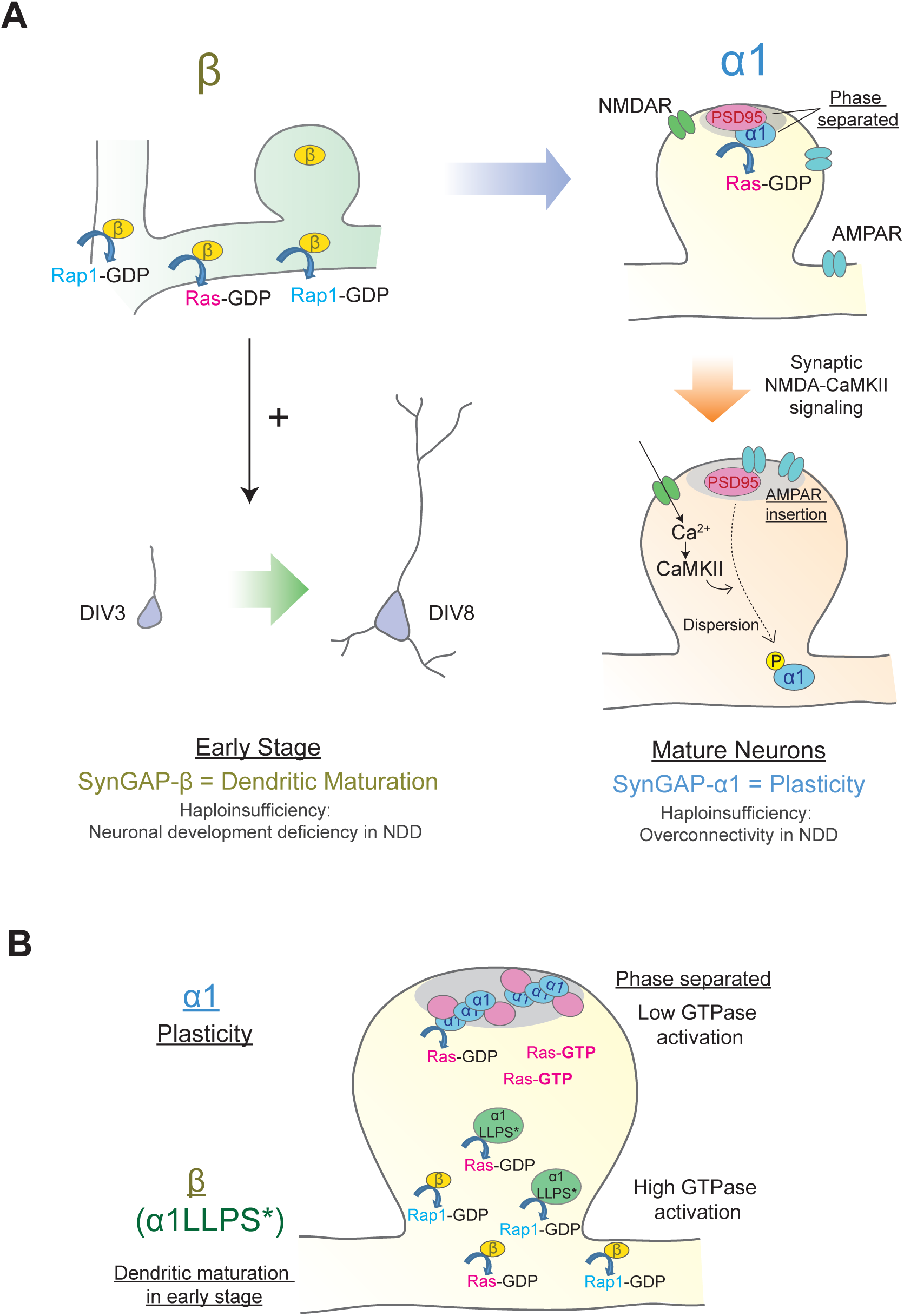
Distinct roles of individual SynGAP isoforms in neuronal development and synaptic plasticity. (A) Schematics illustrating isoform-specific roles for SynGAP in neuronal maturation and synaptic plasticity. SynGAP-β is expressed early in development, has the lowest LLPS propensity resulting in cytosolic localization, possesses the highest GAP activity in cells, and promotes normal dendritic development. SynGAP-β deficiency may be relevant to the neuronal development deficits in neurodevelopmental disorders (NDD). SynGAP-α1 is expressed later in development, undergoes strongest LLPS in spines resulting in dense expression in the PSD at the basal state, and is rapidly dispersed upon synaptic NMDAR-CaMKII activation. SynGAP-α1 deficiency may be relevant to the synaptic plasticity deficits and overconnectivity in NDD. (B) Schematics illustrating the phase-separation and the localization/functions of SynGAP isoforms. SynGAP-α1 is tightly packed at PSD by phase separation and has the ability of low GTPase activations. In contrast, SynGAP-β is less phase-separated and localized more in cytoplasmic region in synapses and dendritic shafts. It has a strong ability of GTPase activation. The phase-separation mutant of SynGAP-α1 LLPS*) behaves similarly to SynGAP-β.

### SynGAP-β: early expression and strongest GAP function - Roles in dendritic development and its implications on neurodevelopmental disorders

In the present study, we discovered that knockdown of SynGAP results in excessive proximal dendritic sprouting in immature hippocampal neurons, and importantly, SynGAP-β effectively rescues this developmental phenotype. Various small G proteins such as Ras, Rap1, Rac1, and RhoA tightly regulate dendritic arbor development by precisely controlling the number and length of dendritic branches (Fu et al., 2007; Nakayama et al., 2000; Saito et al., 2009; Sepulveda et al., 2010). For example, Rap1 increases proximal dendritic branching in rat cortical neurons, and Rap2 activation decreases the length and complexity of developing axonal and dendritic branches (Chen et al., 2005). These data link our observed dendritic phenotype caused by SYNGAP1 deficiency to regulation of small G-proteins. Further, dominant-negative forms of Rac1 decrease proximal dendritic branching and increase distal dendritic branching in hippocampal organotypic slice cultures, suggesting that a proper balance of small G-protein activation is crucial for normal dendritic development (Nakayama et al., 2000). Additionally, the Ras-PI3K–Akt–mTOR pathway controls somatic and dendritic sizes and coordinates with Ras-mitogen-activated protein kinase signaling to maintain dendritic complexity (Kumar et al., 2005). Thus, it is possible that SYNGAP1 haploinsufficiency causes overactivation of Ras and Rap1 and consequently disrupts the balanced signaling required for normal dendritic development. Our results suggest that SynGAP-β plays pivotal role in establishing this balance by regulating Rap1 and other small G proteins. There are currently a variety of available downstream inhibitors of small G proteins, which may prove to be valuable therapeutic targets to decrease dendritic deficits caused by SynGAP-β deficiency.

Recent studies have suggested that human induced pluripotent stem cells (hiPSCs) derived from ASD patients exhibit accelerated dendritic outgrowth and excessive dendritic branching following neuronal differentiation (Schafer et al., 2019). These observations, together with our finding that SynGAP-β predominantly promotes dendritic arbor development, links dendritic morphological deficits to the abnormal neuronal wiring associated with NDDs.

### Critical role of SynGAP-α1 in synaptic plasticity; strong LLPS propensity for synaptic enrichment and dispersion during LTP

We have previously reported that SynGAP-α1 is rapidly dispersed in response to LTP-inducing synaptic activity. This dispersion allows for AMPAR insertion into the synaptic membrane and for enlargement of dendritic spines. Thus, SynGAP-α1 functions to regulate AMPAR accumulation and spine size at basal states to maintain a neuron’s ability to undergo LTP and avoid saturating plasticity (Araki et al., 2015; Zhu et al., 2002). Here, we demonstrate that only the SynGAP-α1 isoform efficiently drives AMPAR insertion and spine enlargements during LTP. SynGAP-α1 is highly concentrated in the PSD via LLPS with PSD-95, which generates a sharp concentration gradient of SynGAP in dendritic spines that is collapsed following activity-dependent SynGAP-α1 dispersion and subsequent synaptic potentiation (Araki et al., 2015; Dosemeci and Jaffe, 2010; Lautz et al., 2018; Lautz et al., 2019; Yang et al., 2013; Yang et al., 2011). We speculate that the high magnitude of activity-dependent dispersion of SynGAP-α1 is due, in part, to the tendency of SynGAP-α1 to robustly phase separate with PSD95 and facilitate LTP-associated signaling in the synapse. Mice with *Syngap1* haploinsufficiency display exaggerated synaptic connectivity and E/I balance in CA1 excitatory neurons (Clement et al., 2012). These may be a result mainly from α1 specific haploinsufficiency as SynGAP-α1 strongly regulates synaptic function compared to other isoforms.

### LLPS propensity of SynGAP isoforms determines its function and GAP activity in neurons

In the present study, an LLPS mutation of the α1 isoform switched its functional outcome from rescuing synaptic plasticity (α1-type) to rescuing dendritic arborization (β type) after SynGAP knockdown, suggesting that distinct structural and LLPS propensities of SynGAP isoforms define roles in regulating neuronal development and/or synaptic plasticity. In addition, we previously showed that SynGAP-α1 LDKD mutant, with its weaker LLPS propensity, is not able to fully rescue knockdown-dependent aberrant synaptic strengthening, and that this rescue results in altered and abnormally lowered LTP threshold (Zeng et al., 2016). Taken together, these results suggest that the LLPS propensity of SynGAP alter its function in neurons and determines between the α1 or β like behavior of the molecules.

As we have shown, the SynGAP isoforms differentially regulate small G-proteins. These results may be due to differences in the localization of the SynGAP isoforms, since rates of biochemical reactions are dependent on the concentration of reactants within a microenvironment. LLPS of SynGAP physically separates its GAP domain within SynGAP from the small G proteins. This is consistent with our observation that while SynGAP-β generally showed the weakest LLPS but has the highest GAP activity towards almost all small G-proteins (**Fig. 7*B***). It is known that various GAPs are also differentially localized by distinct lipid modifications. After synthesis of Ras, Rap1, and Rac1, farnesyl or geranylgeranyl moieties are attached to the C-terminal ‘CAAX’ motifs (C: Cys; A: an aliphatic amino acid, X: M, Q, S, T, or A for farnesyl, L or I for geranylgeranyl) for membrane tethering, facilitating interaction with effector molecules at the proximity of the membrane. The various small G proteins have slightly different CAAX motifs that are susceptible to distinct modifications (For example, H-Ras has CVIM for farnesylation, Rap1a has CLLL for geranylgeranylation) and thus differentially targeted to cellular microenvironments (Moores et al., 1991; Simanshu et al., 2017; Wright and Philips, 2006). Thus, the combinations of small G-protein localization and SynGAP isoform localization may define the ability of each SynGAP isoform to activate distinct GTPase activation, and thus differentiate their function in synaptic spines or dendrites.

### Importance of characterizing various SynGAP isoforms to elucidate ID/ASD pathogenesis: Therapeutic strategies for MRD5 and neurodevelopmental disorders

*SYNGAP1* is the 4th most prevalent gene that is mutated in neurodevelopmental disorders (NDD) such as ID/ASD. Mutation of SYNGAP1 explains ∼0.75% of all NDD cases which is nearly as high as prominent X-linked disorders, such as the Fragile X syndrome (UK-DDD-study, 2015). SYNGAP1 is located in the 6p21 region, near the major immuno-histocompatibility complex (MHC) where a high rate of sequence variability is observed between people (Reche and Reinherz, 2003; Sommer, 2005). This high rate of variability within genomic regions, along with the fact that *SYNGAP1* does not have neuronal homologs despite its critical roles in development and plasticity, may contribute the high rate of association of *SYNGAP1* with MRD5 in ID/ASD populations. We highlight the importance of SynGAP-α1 in synaptic plasticity and suggest that correcting the exaggerated downstream activity due to haploinsufficiency of α1 might be beneficial for the patients. In this paper, we characterized dynamic changes in the expression profile of SynGAP isoforms in the brain throughout development. We showed that SynGAP-α1 expression accounts for only 25∼35% of total SynGAP, highlighting the importance of assessing the function of all isoforms to understand the pathogenesis of MRD5. We also found SynGAP-β to be expressed earliest in development, and to play a unique role in dendritic arbor development. Further characterization of downstream small G proteins and kinases that have pivotal roles in dendritic development will lead to unique targets for treating the aberrant neuronal wiring that is associated with MRD5 as well as other ID/ASD-related neurodevelopmental disorders.

## Materials and Methods

### Reagents

All restriction enzymes were obtained from New England Biolabs. Chemicals were obtained from SIGMA-Aldrich unless otherwise specified. TTX, Bicuculline, and Strychnine were obtained from TOCRIS Bioscience. Goat anti-SynGAP-α1 antibody is from Santa Cruz (sc-8572). Rabbit pan-SynGAP 947-1167 antibody is from Thermo scientific (#PA-1-046). DNA sequencing was performed at the Johns Hopkins University School of Medicine Sequencing Facility.

### Antibodies

The rabbit anti-SynGAP-α1 antibody was used as described in previous reports (Kim et al., 1998; Rumbaugh et al., 2006). To raise antibodies that specifically recognize each non-α1 SynGAP isoform, we conjugated 10-18 amino acids of the C-terminal sequences of each SynGAP isoform with an N-terminal Cysteine (CPPRLQITENGEFRNTADH (JH7265, α2), CGGGGAAPGPPRHG (JH7266, β), and CRLLDAQLLIR (JH7366, γ)) to Keyhole limpet hemocyanin (PIERCE) using the manufacturer’s protocol. Antisera acquired after 2 booster injections (α1, α2, β, and γ) were affinity purified using peptide coupling sulfolink-beads (PIERCE).

### Human SYNGAP1 splicing analysis

Exon junction abundance data was acquired from Brain Seq Consortium Phase 1 (Jaffe et al., 2018). Briefly, total RNA extracted from post-mortem tissue of the dorsolateral prefrontal cortex grey matter (DLPFC) was sequenced and reads were aligned with TopHat (v2.0.4) based on known transcripts of the Ensembl build GRCh37.67. Splice junctions were quantified by the number of supporting reads aligned by Tophat, and counts were converted to “RP80M” values, or “reads per 80 million mapped” using the total number of aligned reads across the autosomal and sex chromosomes (dropping reads mapping to the mitochondria chromosome), which can be interpreted as the number of reads supporting the junction in our average library size, and is equivalent to counts per million reads mapped (CPM) multiplied by 80. For a given 5’ splice donor site, all identified 3’ splice acceptors were grouped together to calculate the relative abundance of each splice decision.

### Liquid-liquid phase separation assay

HEK 293T cells were transfected with SynGAP and/or PSD-95 for 16 h. Cells were lysed in 0.5 ml of assay buffer (50 mM Tris pH 8.0, 100 mM NaCl, 1 mM EDTA, 1 mM EGTA, 1% Triton X-100, 0.1% SDS, 0.5% Sodium deoxycholate, with cOmplete Protease inhibitor EDTA-free mix (Roche/ SIGMA)). Lysates were centrifuged at 15000 x g for 10 min at 4°C. The supernatant-containing the soluble [S] fraction was collected. Pellets were resuspended and sonicated in 0.5 ml of assay buffer to obtain complete homogenate of pellet [P] fraction.

For imaging of LLPS dynamics in living cells, HEK cells were grown on Poly-L-Lysine-coated glass coverslips. Cells were transfected with GFP-SynGAP and/or PSD-95-mCherry for 16 h before being placed in a custom-made live imaging chamber for observation under confocal microscopy. Cells were perfused with extracellular solution (ECS: 143 mM NaCl, 5 mM KCl, 10 mM Hepes pH 7.42, 10 mM Glucose, 2 mM CaCl_2_, 1 mM MgCl_2_). For DAPI staining, cells were fixed with Parafix (4% paraformaldehyde, 4% Sucrose in PBS) for 15 min at room temperature, followed by incubating with 300 nM DAPI in PBS for 5 min at room temperature. Cells were briefly washed with PBS and mounted on slideglass. Cells were observed on an LSM880 (Zeiss) microscopy with a 40x objective lens (NA 1.3).

### PSD fractionation

Fractionation of post-synaptic density (PSD) was performed as previously described (Kohmura et al., 1998). In brief, mouse brains were collected and homogenized by 10-15 strokes of a Dounce A homogenizer in Buffer A (0.32M Sucrose, 10 mM Hepes (pH7.4) with cOmplete protease inhibitor mix (SIGMA)). The homogenate was centrifuged at 1,000 x g for 10 min at 4 °C. The supernatant (Post Nuclear Supernatant; PNS) was collected and centrifuged at 13,800 x g for 20 min at 4 °C. The pellet (P2 fraction) was re-homogenized in 3 volumes of Buffer A. The re-homogenized P2 fraction was layered onto a discontinuous gradient of 0.85, 1.0, 1.2 M sucrose (all containing 10 mM Hepes (pH7.4) plus cOmplete protease inhibitor mix), and were centrifuged at 82,500 x g for 2 h at 4 °C (Beckman SW28 swing rotor). The band between 1.0 and 1.2 M sucrose was collected as the synaptosome fraction and diluted with 80 mM Tris-HCl (pH 8.0). An equal volume of 1 % Triton X-100 was added and rotated for 15 min at 4 °C followed by centrifuging 32,000 x g for 20 min. The supernatant was collected as a Triton-soluble synaptosome (Syn/Tx) fraction, and the pellet was re-homogenized in Buffer A by applying 10 passes through a 21G syringe. Equal amounts of protein (10 μg for immunoblotting) were used for further assay.

### Chemical LTP and quantification

Live imaging and quantification of LTP were performed as described previously(Araki et al., 2015). Hippocampal neurons from embryonic day 18 (E18) rats were seeded on 25-mm poly-L-lysine-coated coverslips. The cells were plated in Neurobasal media (Gibco) containing 50U/ml penicillin, 50mg/ml streptomycin and 2 mM GlutaMax supplemented with 2% B27 (Gibco) and 5% horse serum (Hyclone). At DIV 6, cells were thereafter maintained in glia-conditioned NM1 (Neurobasal media with 2mM GlutaMax, 1% FBS, 2% B27, 1 x FDU (5mM Uridine (SIGMA F0503), 5 mM 5-Fluro-2’-deoxyuridine (SIGMA U3003). Cells were transfected at DIV17-19 with Lipofectamine 2000 (Invitrogen) in accordance with the manufacturer’s manual. After 2 days, coverslips were placed on a custom perfusion chamber with basal ECS (143 mM NaCl, 5 mM KCl, 10 mM Hepes pH 7.42, 10 mM Glucose, 2 mM CaCl_2_, 1 mM MgCl_2_, 0.5 μM TTX, 1 μM Strychnine, 20 μM Bicuculline), and time-lapse images were acquired with either LSM510 (Carl Zeiss; Fig. 1, 4, and 6) or Spinning disk confocal microscopes controlled by axiovision software (Carl Zeiss; Fig. 2, 3, and 5). Following 5-10 min of baseline recording, cells were perfused with 10 ml of glycine/ 0Mg ECS (143 mM NaCl, 5 mM KCl, 10 mM HEPES pH 7.42, 10 mM Glucose, 2 mM CaCl_2_, 0 mM MgCl_2_, 0.5 μM TTX, 1 μM Strychnine, 20 μM Bicuculline, 200 μM Glycine) for 10 min, followed by 10 ml of basal ECS. To stabilize the imaging focal plane for long-term experiments, we employed Definite focus (Zeiss). For quantification, we selected pyramidal neurons based on morphology that consisted of a clear primary dendrite, and quantified all spines on the 30-40 μm stretch of the secondary dendrite beginning just after the branch from the primary dendrite. For identifying spine regions, we used the mCherry channel to select the spine region that was well separated from dendritic shaft. These regions of interest (ROIs) in the mCherry channel were transferred to the green channel to quantify total SynGAP content in spines. Total spine volume was calculated as follows; (Average Red signal at ROI – Average Red signal at Background region) * (Area of ROI). Total SynGAP content was calculated as follows; (Average Green signal at ROI – Average Green signal at Background region) * (Area of ROI). Through this quantification, we can precisely quantify the total signals at each spine even if the circled region contained some background area. For [%] spine enlargement before/ after LTP, we took a relative ratio of these total spine volume (total red signal) of each spine before/ after LTP ([%] spine enlargement = (Total Red Signal after chemLTP / Total Red signal at basal state-1)*100). For [%] SynGAP dispersion, we calculated the degree of total SynGAP content loss after chemLTP at each spine compared to the total SynGAP content at basal state ([%] dispersion = (1-Total Green Signal after chemLTP / Total Green signal at basal state) * 100).

### Dendritic arbor development assay

Cultured hippocampal neurons were plated on coverslips as described above and were co-transfected at DIV 3-4 with pSUPER-SynGAP shRNA and shRNA-resistant GFP-SynGAP-α1, α2, β, and γ replacement constructs. pCAG-DsRed2 was also co-transfected as a cell-fill for morphological analysis. Neurons were fixed at DIV 8-9 by incubating them with Parafix (4% paraformaldehyde, 4% Sucrose in PBS) for 15 min at room temperature, followed by incubation with 300 nM DAPI in PBS for 5 min at room temperature. Cells were briefly washed with PBS and mounted onto glass slides. Cells were imaged with a LSM880 (Zeiss) confocal microscope equipped with a 40x objective lens (NA 1.3) and GaAsP detectors. To obtain Sholl profiles of dendritic arbors (Sholl, 1953), images of entire dendritic arbors of hippocampal neurons expressing DsRed were acquired and processed using Image J (Fiji) software. Scholl analysis consisted of drawing concentric rings with radii of 10, 20, 30, 40, 50, 100, and 150 μm from the center of the cell body and counting the number of dendritic intersections across each concentric circle. If a branch point fell on a line, it was counted as two crossings (Nakayama et al., 2000).

### Small GTPase activity assay

Small GTPase activity was measured using a small GTPase-GTP pull-down assay. DNA constructs expressing a small G protein and a single SynGAP isoform were co-transfected into HEK cells for 48-72 hours. Active Ras levels were then assayed using a Ras activation assay kit (EMD Millipore). In brief, cells were lysed in Mg^2+^ lysis/wash buffer (25 mM HEPES pH 7.5, 150 mM NaCl, 1% Igepal CA-630, 10 mM MgCl_2_, 1 mM EDTA, 10% glycerol), and active GTP-bound small G-proteins were pulled down using beads covalently bound to effector domains. After washing beads, active GTP-bound small G proteins were recovered through the addition of 2x SDS sample buffer followed by SDS-PAGE and subsequent immunoblotting for the various small G proteins.

### Statistics

All data are expressed as means ± S.E.M. of values. One-way ANOVAs were used, followed by Tukey post hoc for multiple comparisons unless otherwise specified. If the interaction between two-factors was observed by two-way ANOVA, we performed individual Tukey *post hoc* tests to compare the measures as a function of one factor in each fixed levels of another factor unless otherwise specified. Statistical analyses and preparations of graphs were performed using SPSS 9.0, Excel 2010, or GraphPad Prism 4.0 / 5.0 software (\**p* < 0.05; \*\**p* < 0.01; \*\*\**p* < 0.001).

## Author contributions

Y.A. and R.H. designed experiments. Y.A., I.H., T.G., and S.J. performed experiments. Y.A., I.H., T.G., and R.H. contributed new reagents / analytic tools. Y.A., I.H., and R.H. analyzed data. Y.A., T.G., and R.H. wrote the paper.

## Acknowledgements

We thank all members of the Huganir lab for discussion and support throughout this work especially Drs. Kacey Rajkovich, Elizabeth Gerber, Megnan Tian, and Bian Liu for critical reading and preparation of the manuscript. We also thank Andrew E. Jaffe, and Daniel R. Weinberger for help with acquiring and analyzing of RNAseq data. This work was supported by grants from National Institute of Health (MH112151, NS036715) and the SynGAP Research Fund. We want to thank the Bridge The Gap SYNGAP Education and Research Foundation, the SynGAP Research Fund, and all of the SynGAP patient families for their outreach and advocacy.

