## Supplementary Information for "SynGAP splice isoforms differentially regulate synaptic plasticity and dendritic development"

**2[+]. Lieber Institute for Brain Development**

#### Supplementary figure legends

**Supplementary Fig. 1 (Supplementary information of Fig. 1) SynGAP isoforms are differentially expressed during brain development**

(A) Quantification of Fig. 1C immunoblot showing specificity of SynGAP isoform-specific antibodies. Graph of relative SynGAP isoform levels with respect to total SynGAP expression measured from immunoblot was displayed. Error bars indicate ± SEM. Two-way ANOVA followed by Tukey's post hoc test (Isoform $F(4,30) = 1.900$; $p = 0.13$, Genotype $F(1,30) = 451.2$; $p < 0.001$, Interaction $F(4,30)=1.900$; $p = 0.13$, $n = 4$ each condition) was performed. Error bar indicates ± SEM.

(B) Quantification of Fig. 1D immunoblot showing endogenous expression and distribution of SynGAP isoforms in various organs. Heat map of immunoblot was displayed. The amount of protein in the brain is standardized as 1.0.

(C) Quantification of Fig. 1E immunoblot showing western blot of endogenous levels of individual SynGAP isoforms and other synaptic proteins in lysates prepared from several brain regions obtained from *WT*. (OB: Olfactory bulb, CC: Cerebral cortex, Hip: Hippocampus, ST: Striatum, Th: Thalamus, Mid: Midbrain, Ce: Cerebellum). Two-way ANOVA followed by Tukey's post hoc test (Brain regions $F(7, 264) = 1048$ ; $p < 0.0001$, Molecules $F(10,264) = 8.0 \times 10^{-12}$ ; $p > 0.9999$, Interaction $F(70.264) = 59.06$ ; $p<0.0001$, $n = 4$ each condition) was performed. The mean values of each signals were plotted in the graph.

(D) Complete set of quantification for immunoblots representing relative enrichments along developmental stage corresponds to Fig. 1G. The mean values of each signals were plotted in the graph. Error bars indicate ± SEM.

(E) mRNA expression of the β and non-β SYNGAP1 isoforms across age in human dorsolateral prefrontal cortex. RNAseq reads spanning the exon 17-18 junction supporting either isoform were normalized to sequencing depth and plotted against human age (post-conception weeks and years) with a linear regression.

(F) mRNA expression of the α1, α2, and γ SYNGAP1 isoforms across age. RNAseq reads spanning the exon 18-19 junction (γ) or exon 18-20 junction (α1, α2) supporting each isoform were normalized to sequencing depth and plotted against human age.

**Supplementary Fig. 2 (Supplementary information of Fig. 2) Phase separation and subcellular localization of various SynGAP isoforms**

(A) Representative immunoblot probing levels of GFP-SynGAP (WT or phase separation mutant) and myc-PSD95 in phase-separated supernatant and pellet lysate fractions obtained from HEK cells expressing myc-PSD95 and either GFP-SynGAP-WT or GFP-SynGAP-LDKD constructs. (Right panel) Quantification of pellet fraction ratios obtained from averaged immunoblots as the representative example shown in (A). Error bars indicate ± SEM. Two-way ANOVA followed by Tukey's post hoc test (Molecules $F(1,30) = 3.026$; $p = 0.09$, Transfections $F(4,30) = 280.7$; $p < 0.0001$, Interaction $F(4,30)=59.69$ ; $p < 0.0001$, $n = 4$, *** $p<0.001$, ** $p<0.01$, * $< 0.05$) was performed.

(B) Representative immunoblot probing levels of GFP-SynGAP (WT or PDZ deletion mutant) and myc-PSD95 in phase-separated supernatant and pellet lysate fractions obtained from HEK cells expressing myc-PSD95 and either GFP-SynGAP-WT or GFP-SynGAP-ΔQTRV (PDZ ligand deletion) constructs. (Right panel) Quantification of pellet fraction ratios obtained from averaged immunoblots as the representative example shown in (B). Error bars indicate ± SEM. Two-way ANOVA followed by Tukey's post hoc test (Molecules $F(1,24) = 195.00$; $p < 0.0001$, Transfections $F(3,24) = 456.7$; $p < 0.0001$, Interaction $F(3,24)=71.27$ ; $p < 0.0001$, $n = 4$, *** $p<0.001$, ** $p<0.01$, * $< 0.05$) was performed.

(C) Complete set of immunoblot in Fig. 2B showing representative western blot probing levels of individual SynGAP isoforms in phase-separated supernatant and pellet lysate fractions obtained from HEK cells expressing myc-PSD95 and individual GFP-tagged SynGAP isoforms.

(D) Representative confocal images of fixed HEK cells expressing myc-PSD95 alone or myc-PSD95 and either GFP-SynGAP-WT or GFP-SynGAP-LDKD constructs (F1). Scale Bar, 10 μm. Arrowheads indicate PSD95- and SynGAP-α1-containing puncta (>1 μm). (F2) Quantification of the averaged percentage of PSD95-positive puncta identified in images of fixed HEK cells as shown in (F1). Error bars indicate ± SEM. One-way ANOVA followed by Tukey's post hoc test (Transfections $F(2, 9) = 126.8$; $p < 0.0001$, n = 4 independent coverslip, *** $p<0.001$, ** $p<0.01$, * < 0.05) was performed.

(E) Complete set of quantification for immunoblot probing endogenous levels of individual SynGAP isoforms and other synaptic proteins in forebrain tissue lysates obtained from adult mice subjected to postsynaptic density fractionation that corresponds to Fig. 2G. Error bars indicate ± SEM.

**Supplementary Fig. 3 (Supplementary information of Fig. 4-6) shRNA efficiency and rescue SynGAP construct titration assay**

(A) Efficiency of shRNA-SynGAP construct. Hippocampal neurons were transfected with shRNA-SynGAP#5 construct together with GFP (marker for transfected cell) and stained for pan-SynGAP antibody (** $p < 0.01$, 77.3 ± 0.1 % reduction of SynGAP expression upon our shRNA-SynGAP#5 construct, n = 4 cells, unpaired T-test (two-tailed), Error bars indicate ± SEM). Scale Bar, 10 μm.

(B) Expression levels of our shRNA-resistant SynGAP isoform construct. Hippocampal neurons were transfected with mCherry and GFP-SynGAP isoform construct and stained for SynGAP isoform specific antibodies described in Fig. 1 (α1 88.3 ± 0.2 %, α2 97.0 ± 0.2%, β 83.5 ± 0.1 % for GFP-SynGAP portion, n = 4 cells). Dotted line showed endogenous SynGAP expression levels. Error bars indicate ± SEM. One-way ANOVA followed by Tukey's post hoc test (Transfections $F(4,10) = 126.8$; $p < 0.01$, n = 3 cells, * $p < 0.05$) was performed. Scale Bar, 10 μm.

Supplementary Fig. 1

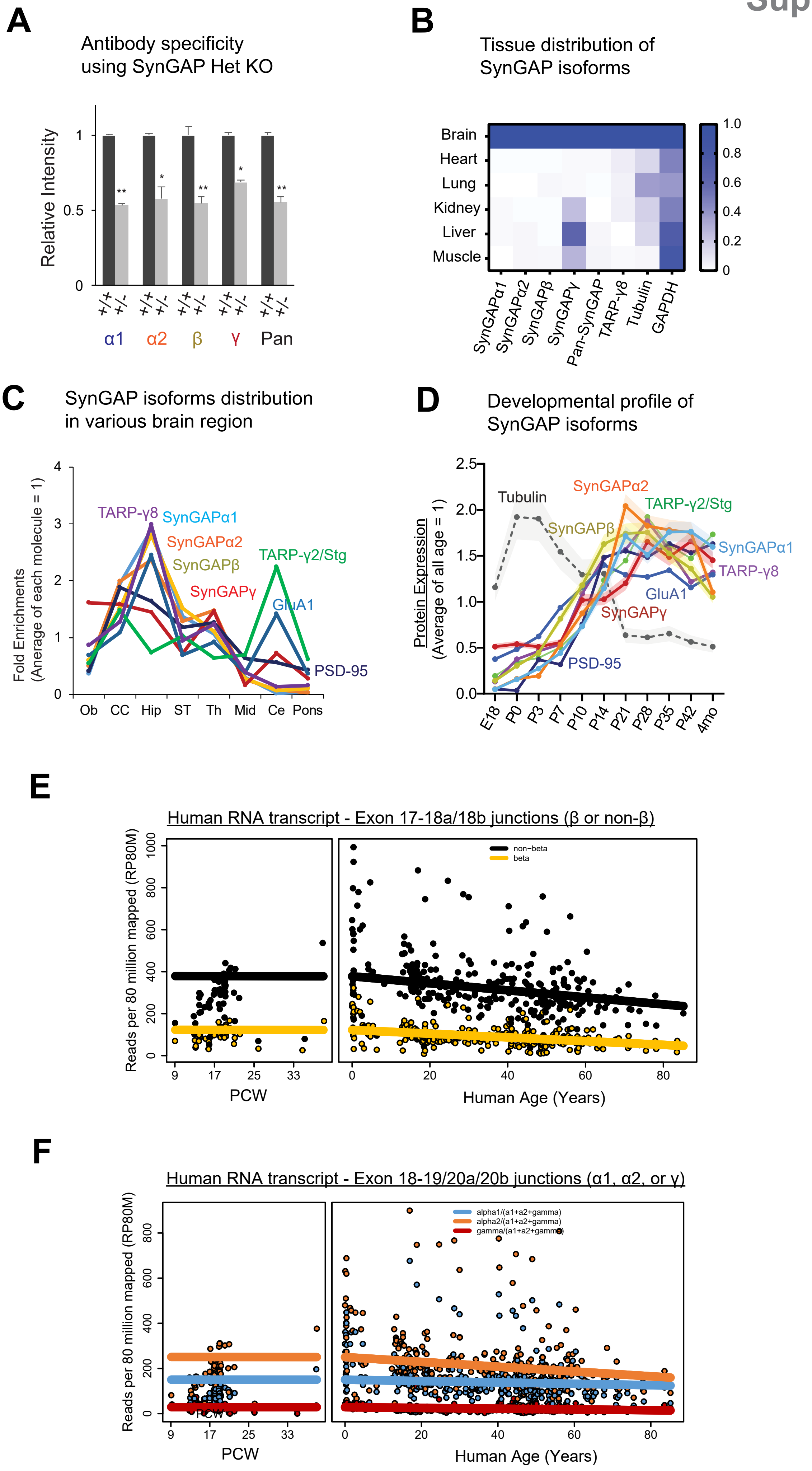

Supplementary Fig. 2 (1/2)

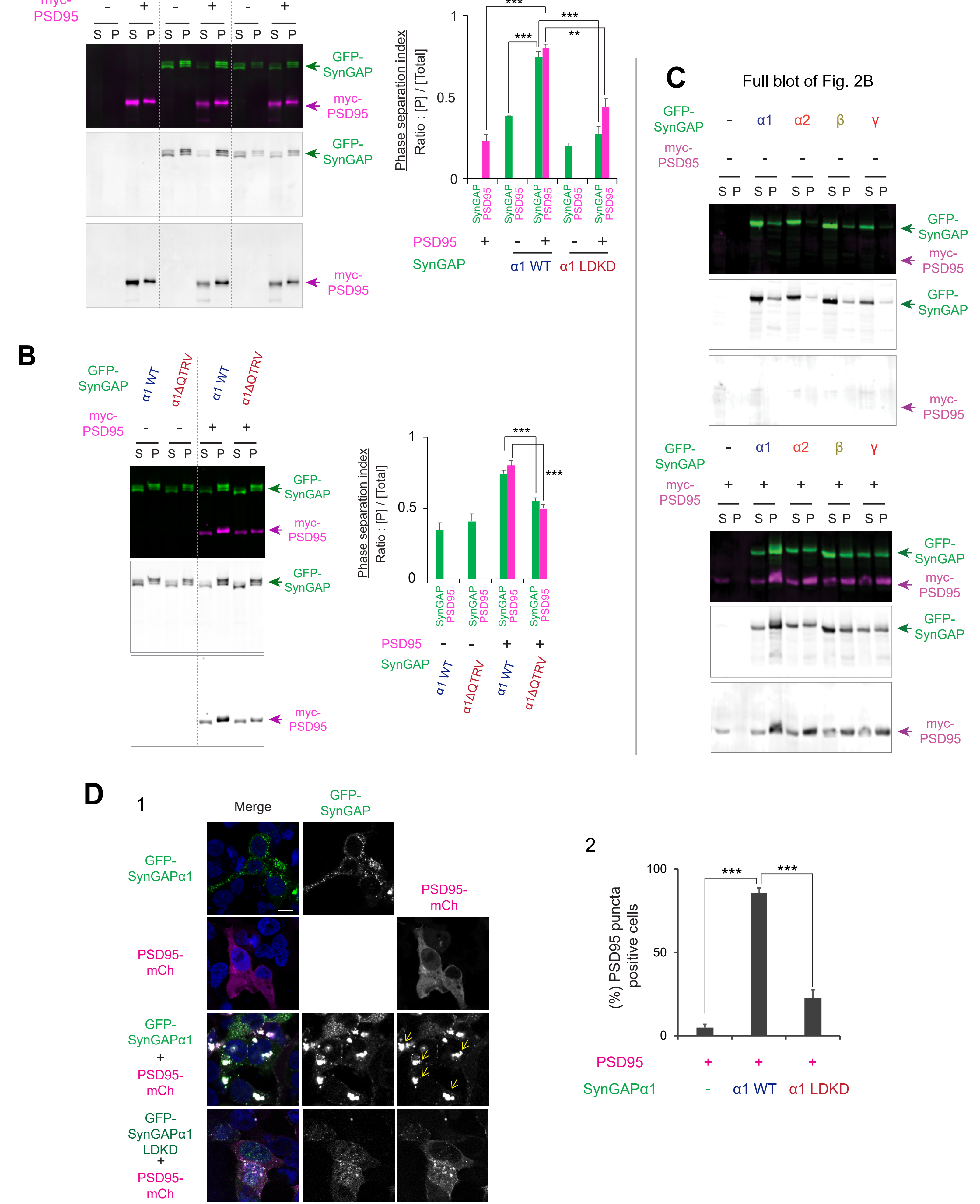

**E**

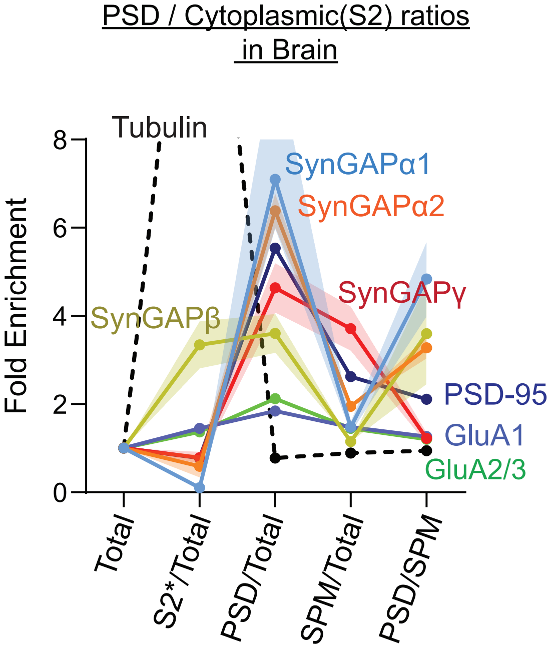
PSD / Cytoplasmic(S2) ratios
in Brain
Fold Enrichment
8
6
4
2
0
Tubulin
SynGAPα1
SynGAPα2
SynGAPγ
SynGAPβ
PSD-95
GluA1
GluA2/3
Total
S2*/Total
PSD/Total
SPM/Total
PSD/SPM

### Supplementary Fig. 3

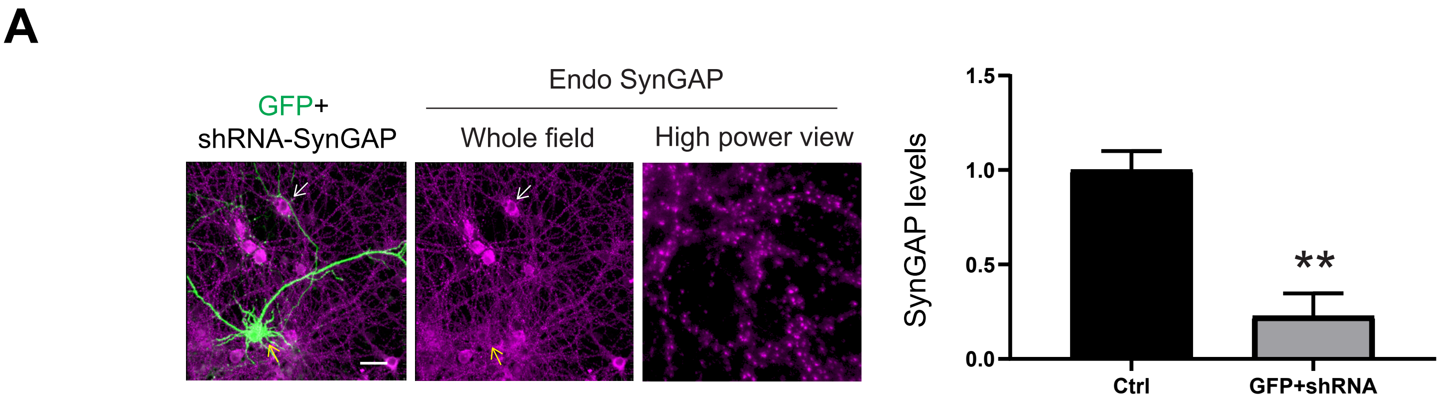


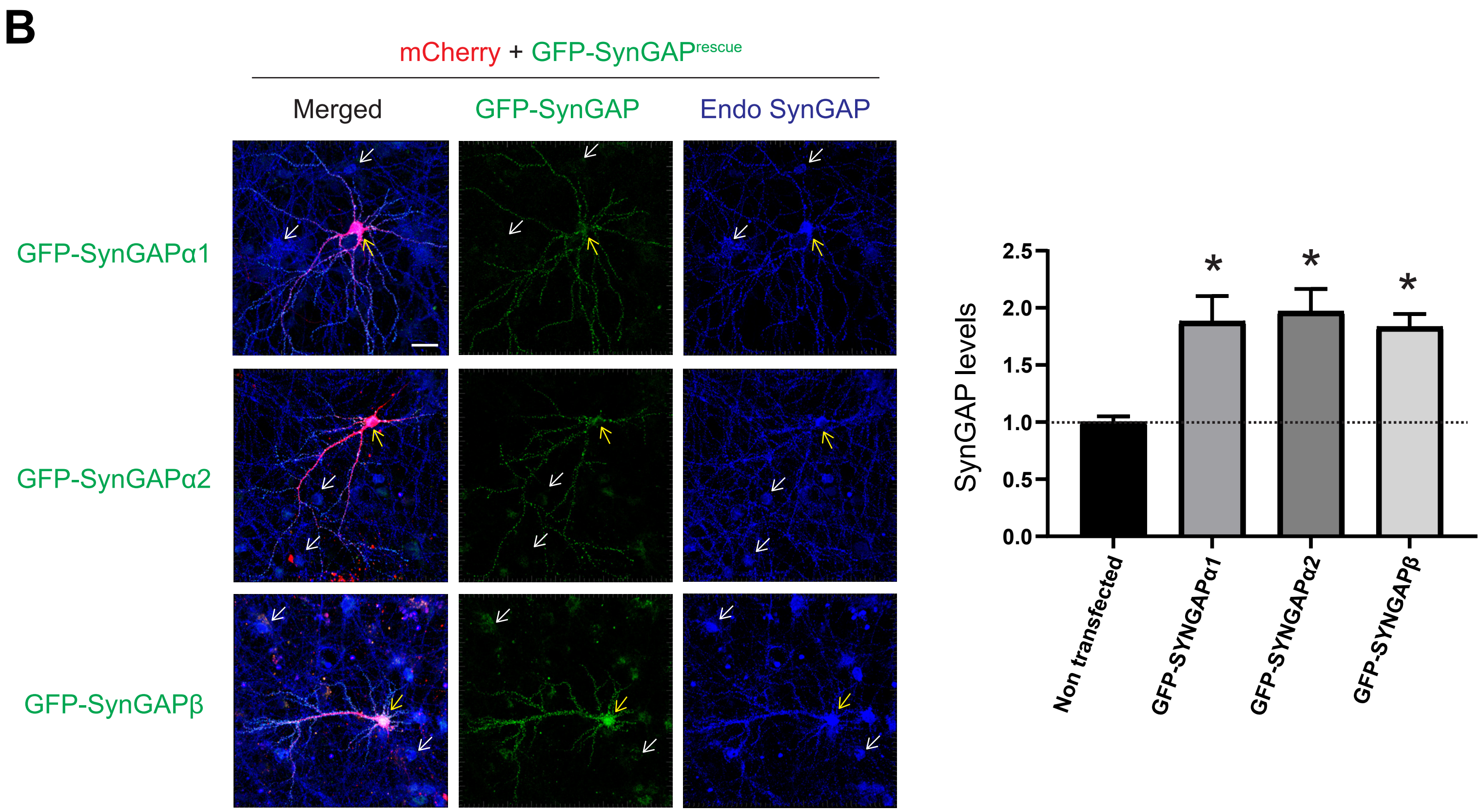
